# Integrated Protocol of Protein Structure Modeling for Cryo-EM with Deep Learning and Structure Prediction

**DOI:** 10.1101/2023.10.19.563151

**Authors:** Genki Terashi, Xiao Wang, Devashish Prasad, Tsukasa Nakamura, Han Zhu, Daisuke Kihara

**Affiliations:** Department of Biological Sciences, Purdue University, West Lafayette, Indiana, 47907, USA; Department of Computer Science, Purdue University, West Lafayette, Indiana, 47907, USA

**Keywords:** electron microscopy, cryo-EM, protein structure modeling, deep learning, structural biology, protein structure prediction

## Abstract

Structure modeling from maps is an indispensable step for studying proteins and their complexes with cryogenic electron microscopy (cryo-EM). Although the resolution of determined cryo-EM maps has generally improved, there are still many cases where tracing protein main-chains is difficult, even in maps determined at a near atomic resolution. Here, we have developed a protein structure modeling method, called DeepMainmast, which employs deep learning to capture the local map features of amino acids and atoms to assist main-chain tracing. Moreover, since Alphafold2 demonstrates high accuracy in protein structure prediction, we have integrated complementary strengths of de novo density tracing using deep learning with Alphafold2’s structure modeling to achieve even higher accuracy than each method alone. Additionally, the protocol is able to accurately assign chain identity to the structure models of homo-multimers.

## Main Text

An increasing numbers of protein structures have been modeled from cryo-electron microscopy (cryo-EM) maps. This trend is expected to further grow as the improvements in cryo-EM techniques have overcome conventional limitations in terms of the resolution, the molecular size, and types of proteins to be determined^1^. Although map resolution has generally improved steadily over the past years, there are still many situations in practice where modelers face difficulties in tracing main-chains due to locally low resolution in the map, among other factors. Moreover, it is observed that modeling errors frequently occur, even in maps determined at around subatomic resolution^2^.

To aid in structure modeling from cryo-EM maps, a new generation of modeling methods has been developed over the past few years^3^. These methods include those that trace the main-chain through local points with high density values in the map using graph-based approaches^4–6^, a method that uses structure fragment library^7,8^ or structural prediction^9^. Additionally, some methods use deep learning to identify local map patterns that correspond to amino acids^10,11^. These recent methods are particularly useful for de novo structure modeling, where experimentally determined structures are not available for fitting in maps with a resolution of around 3 to 5 Å, which is beyond the scope of conventional modeling tools for X-ray crystallography^12,13^.

When discussing about protein structure modeling of any kind, it is essential to keep in mind the recent advances in protein structure prediction. Since the appearance of Alphafold2 (AF2), a highly accurate protein structure prediction method^14^, structure modeling for cryo-EM has also undergone changed, as structures predicted by AF2 could be simply fit to the map if it is expected to be at a high accuracy^15^. There are numerous reported cases where models generated by AF2 were accurate enough to be fitted in the map^16^. However, on the other hand, there are also many instances where AF2 models do not agree with the map.

Considering all the factors surrounding structure modeling for cryo-EM, here we developed an integrated protein structure modeling protocol that models protein structures for a cryo-EM density map using deep learning in the map and also considers AF2 models when applicable. The primary component of the protocol is DeepMainmast(base), a new de novo protein main-chain tracing method guided by identified positions of Cα atoms and types of amino acids by deep learning. Compared to its predecessor, MAINMAST^5^, our new method features several improvements, including the use of deep learning and a more effective main-chain tracing approach called the Vehicle Routing Problem^17^ (VRP) solver^18^, which is better suited for multi-chain complexes. In addition to generating structure models with DeepMainmast(base), we also employ AF2 and blend its model with those from DeepMainmast(base) when they exhibit consistency in local structures. The protocol generates a total of 20 models, with different levels of blending of models, which are finally ranked based on the DOT score, a map-model fitness score, and the DAQ score, a model quality assessment score we recently developed^2^. The entire protocol, DeepMainmast, was shown to substantially outperform existing methods including AF2 on datasets of experimental EM maps of single-chain proteins and multi-chain protein complexes in terms of the main-chain conformation and sequence assignment accuracy. The protocol is also equipped with the ability to accurately connect individual chains of homo-multimers, which is not an easy task and has not been addressed in other methods.

## Workflow of the Modeling Protocol

The protein modeling protocol of DeepMainmast takes a cryo-EM map and sequences of proteins that are included in the map as input and outputs protein structure models. The models are sorted by the scoring function that indicates fit to the map. The overall protocol is shown in Fig. 1. The main and foremost component of the pipeline is DeepMainmast(base), the new deep learning-assisted protein structure modeling method. In parallel to the model building with DeepMainmast(base), protein structures will be also predicted from the input sequences by AF2, which are hybridized with DeepMainmast(base) models if local conformations are compatible with each other. We also fit the entire AF2 models to the map using our VESPER algorithm, which finds the optimal superimposition by considering local tensors of density^19^. Models fitted to the map are finally ranked by the scoring function, which combines DAQ^2^ and DOT^19^, scores that evaluate amino acids and atom propensity of local maps and local tensor agreement used in VESPER, respectively. In what follows, we briefly explain each step. For more details see Methods.

**Figure 1.**
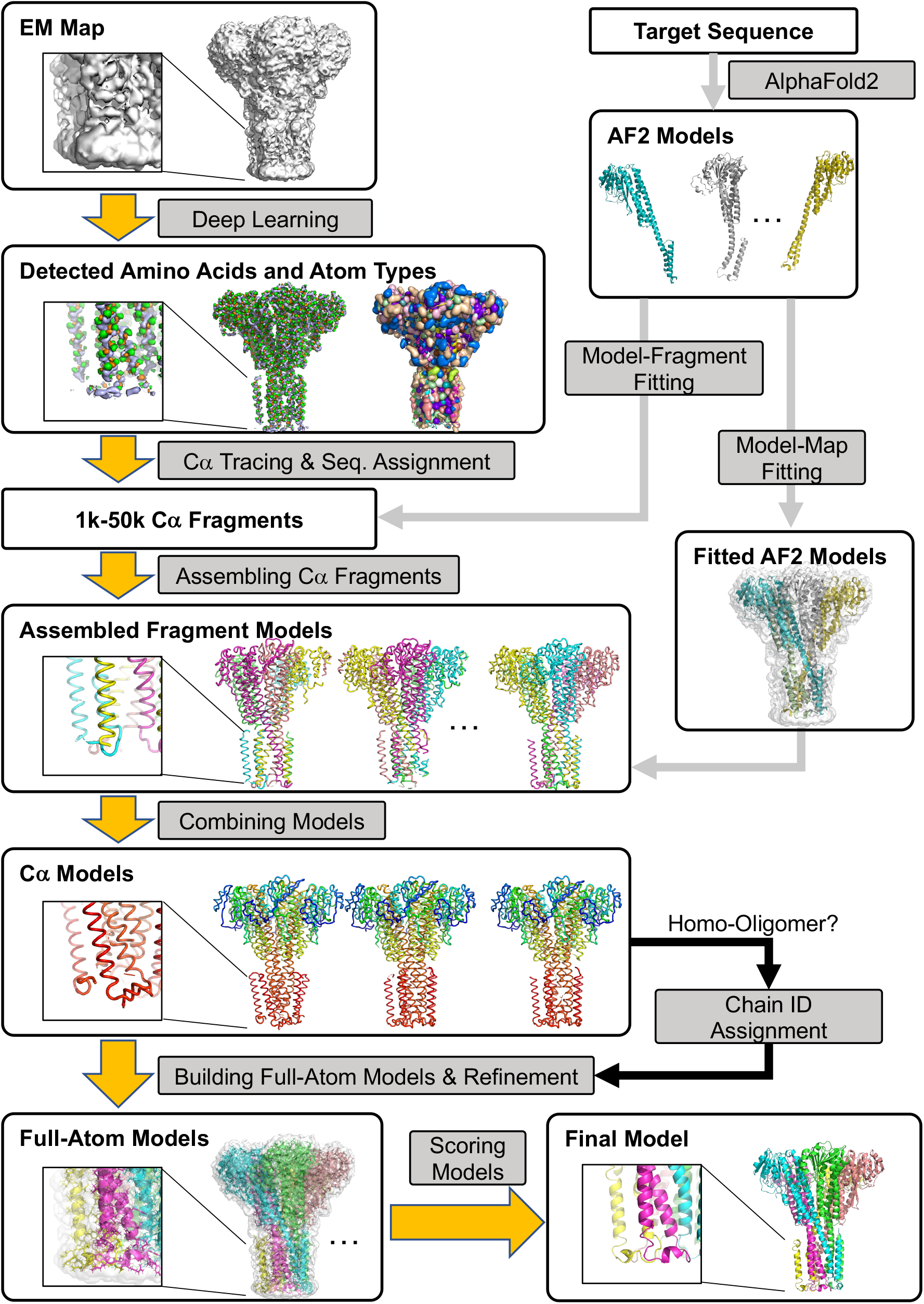
Overview of DeepMainmast protocol. We used a homo-multimer complex, magnesium channel CorA (PDB ID: 3JCF, EMID-6551, resolution: 3.8 Å), as an example. The magnified images in boxes highlight the transmembrane region. The yellow arrows represent the core of the protocol, DeepMainmast(base). It consists of six logical steps: (1) Detecting amino-acid types and atom types using deep learning (Emap2sf). The image on the left shows the detected atom types (Cα atom: green, carbon: orange, and nitrogen: light blue). The image on the right shows the detected amino acid types in different colors. (2) Tracing Cα path and assigning the target sequence using the Vehicle Routing Problem Solver and the Dynamic Programming algorithm. Different parameter combinations are used. (3) Assembling Cα fragments using the Constraint Problem (CP) Solver. Colors indicate chain IDs. (4) Combining Cα models built under different parameter combinations using the CP Solver. Colors indicate the direction of chains from blue to red for the N-terminal to the C-terminal residues. (5) Full-atom building and refinement using PULCHRA and Rosetta-CM. (7) Scoring generated full-atom models based on the DAQ(AA) score and the DOT score. For homo-oligomer targets, chain IDs are assigned based on the structural similarity of homomer proteins (black arrows). The gray arrows represent the procedure to use AF2 models. AF2 models were integrated into the modeling process in two ways, (i) Imposing fragments in the AF2 models to Cα fragments; and by (ii) Fitting AF2 models to the EM map using Map-Model fitting program, VESPER.

## DeepMainmast(base)

### Predicting amino acid types, and atom types using deep learning

The first step of the main-chain modeling by DeepMainmast(base) is to detect amino acids and atoms in a given EM map using a deep learning-based method, Emap2sf. Emap2sf uses a network of the U-net architecture^20^ and computes probabilities of twenty amino-acid types and six atom types (N, Cα, C, O, Cβ, Others) for each grid point in the density map. Computed probability values of twenty amino-acid types and three atoms (N, Cα, C) that form the backbone of protein structure are used in the following Cα tracing step.

### Tracing Cα paths and assigning the protein sequence

Grid points that have a high probability for Cα are clustered by the mean shift algorithm^21^ to generate representative points, named Local Dense Points (LDPs). Then, LDPs are connected to produce Cα paths using a Vehicle Routing Problem (VRP^17^) Solver. VRP is a variation of the traveling salesman problem except that it uses multiple “vehicles” rather than a single “salesman” to explore the node space and connects nodes. Each vehicle in the VRP Solver explores the optimal routes from a pseudo starting point to connect a set of LDPs with minimum total costs of routes under specified constraints. We defined the cost between two LDPs based on the distance and the lowest probability of main-chain atoms along the path of two LDPs.

Once Cα paths are computed, they are aligned with the target sequence using the Smith-Waterman Dynamic Programming (DP) algorithm. To define the matching score between a Cα position in the path and each amino-acid type, we used the DAQ(AA) score^2^ computed from the Emap2sf output to define the matching score between a Cα position in the path to each amino-acid type. DAQ(AA) score was originally designed in our previous work^2^ to evaluate the compatibility of amino acids in a protein model to corresponding local density in a cryo-EM map and here we use the score for modeling. Typically, 1,000-50,000 Cα path-sequence alignments, which we refer to as Cα fragments, are generated for a single-chain protein. We perform this process for each of combinations of three parameters: the Cα probability cutoff, the number of vehicles, and a parameter defining the cost function (see Methods).

### Integration of AF2 structure prediction models

In the DeepMainmast protocol, we integrate predicted protein structures by AF2 (the right branch in Fig. 1). Although AF2 models are not always correct, they can complement structure models from DeepMainmast(base) that result from the density tracing of the EM density. Particularly, AF2 models may be useful for building models in local regions of the map that have low resolution where detecting backbone atoms is not successful. The AF2 model is integrated into DeepMainmast protocol in two steps; as a source of Cα fragments and as a global structure that is fit to the map (two gray arrows in Fig. 1 from AF2 models). We used AF2 version 2.1.0 and ran it without template protein structure data. Out of five full-atom models that AF2 generates, we used the top-ranked model based on the predicted local quality, pLDDT score.

### Enlarging the Cα fragment library with fragments from the AF2 model

The AF2 model was superimposed on the Cα fragments, and structure regions consistent with the Cα fragments were extracted and added to the Cα fragment library. Local structures from AF2 model are extracted only when it agrees with a Cα fragment traced from the density to ensure structural consistency. The main benefit of this process is to supplement the Cα fragment library with AF2 fragments that fill missing regions, such as loop regions or terminal regions in the protein, which correspond to low-density regions in the map and could not be traced. Up to this point in the protocol, four parameters were used. We constructed a Cα fragment library for each of the parameter combinations, which resulted in total of 54 libraries for single-chain proteins and 108 libraries for multi-chain protein complexes. In the next step, a Cα protein model is generated for each of the libraries.

### Assembling Cα fragments to build protein models

Generated Cα fragments are assembled to build Cα structure models. For each of the libraries, one model is generated. This process is considered as an optimization problem of selecting fragments, which we solve with the Constraint Programming (CP)^18^ solver. The CP solver explores feasible combinations of Cα fragments that maximize the total DAQ score while keeping the consistencies of combined fragments. Three types of constraints are considered in the process: (1) no steric collision between Cα fragments; (2) no inconsistent positions for the same amino acid from different fragments; if the same amino acid exists in different fragments, the locations of the amino acids should not be more than 3.5 Å apart; (3) no inconsistent Cα−Cα distance between Cα atoms from different fragments.

### Fitting the AF2 model to the EM density map by VESPER

In addition to assembling Cα fragments for each of the libraries, we also simply superimpose the AF2 model to the density map with the structure fitting program, VESPER^19^, as candidates of structure models (the second gray arrow from the AF2 model box in Fig. 1). Density maps are simulated from the AF2 model at three different resolutions and ten superimpositions are computed for each of them. This gives in total of 30 superimposed AF2 model on the density map.

### Combining assembled Cα fragment models

Up to this point, 54 or 108 assembled Cα fragment models for a single-chain and a multi-chain complex target, respectively, and 30 superimpositions of the AF2 model are generated. Since the fragment-based models may still have gaps and errors in sequence assignment in local regions, we further combine Cα fragment models. Each of the Cα models is fragmented into ten-residue-long Cα fragments, and then the new Cα fragments are assembled into new Cα models by the CP Solver. The same assembling method using the CP Solver is used as described above. The 54 or 108 assembled Cα fragment models are classified into three groups depending on the origin of the fragments, and one final Cα model is constructed from each group (see Methods). 30 superimpositions of the AF2 model are separately considered, and one final model is constructed by assembling them. Thus, in total, four Cα models are generated.

### Chain ID assignment for homo-multimer targets

When modeling a target with multiple chains, the chain ID is naturally labeled to part of the structure model when the sequence is assigned. However, assigning chain ID is not straightforward when the target has multiple chains with the same sequence. To address this issue, we developed a specific step that optimizes consistent chain ID assignment for cases when the target is homo-multimer (Method). This step is unique in DeepMainmast as no other existing method has a similar implementation.

### Building full-atom models

The generated Cα models using the CP Solver are then subjected to full-atom structure building using PULCHRA^22^. Missing regions in all full-atom models are subsequently filled and refined by the Rosetta-CM protocol^7^ with the target sequence and the EM density map. For each Cα model, Rosetta-CM generates five full-atom models, resulting in total of twenty models generated from the four Cα models.

### Evaluating generated models

The twenty full-atom models are evaluated and ranked with two evaluation scores, DAQ score^2^ and DOT score^19^. The DAQ(AA) score evaluates the fit of the amino acid assignment at each map position, while the DOT score from our VESPER program evaluates the agreement of local gradient directions of densities of the map and the simulated map from the protein model. The final score of the full-atom model is the sum of the normalized DAQ score and DOT score.

## Results

### Evaluation of DeepMainmast(base) on single-chain modeling

First, we evaluated the performance of DeepMainmast(base), the *de novo* model building part without the integration of AF2 (yellow arrows in Fig. 1). We used a dataset of single-chain structures from 29 experimental EM maps used in a previous study^5^ (Supplementary Table 1) so that we can compare with two existing methods that used this dataset. In this dataset, the deposited PDB structures were used as the native (reference) structure to be compared against structure models. The density regions that correspond to proteins in the EM maps were manually segmented from the entire EM map using the “zone” tool in UCSF Chimera^23^.

We discuss two versions of the models from DeepMainmast(base). The first version is models that were built by the procedure applied up to the Cα model building (Fig. 1) without applying the subsequent full-atom building step. The second version of the models are those which were built after the full atom building step. We show results of these two versions but mainly discuss the first version in Fig. 2 because the full-atom building step uses Rosetta-CM^7^, which fills gaps in a structure model that may distract objective comparison with existing methods. As the baseline, results were compared with two other *de novo* modeling methods, MAINMAST^5^ and DeepTracer^10^, which used the same dataset in their works. MAINMAST is the original version of the current DeepMainmast, which simply traces locally dense points in a cryo-EM map without using deep learning. DeepTracer has a similar architecture as DeepMainmast(base) as it uses deep-learning to detect main-chain positions and amino-acid types.

**Figure. 2.**
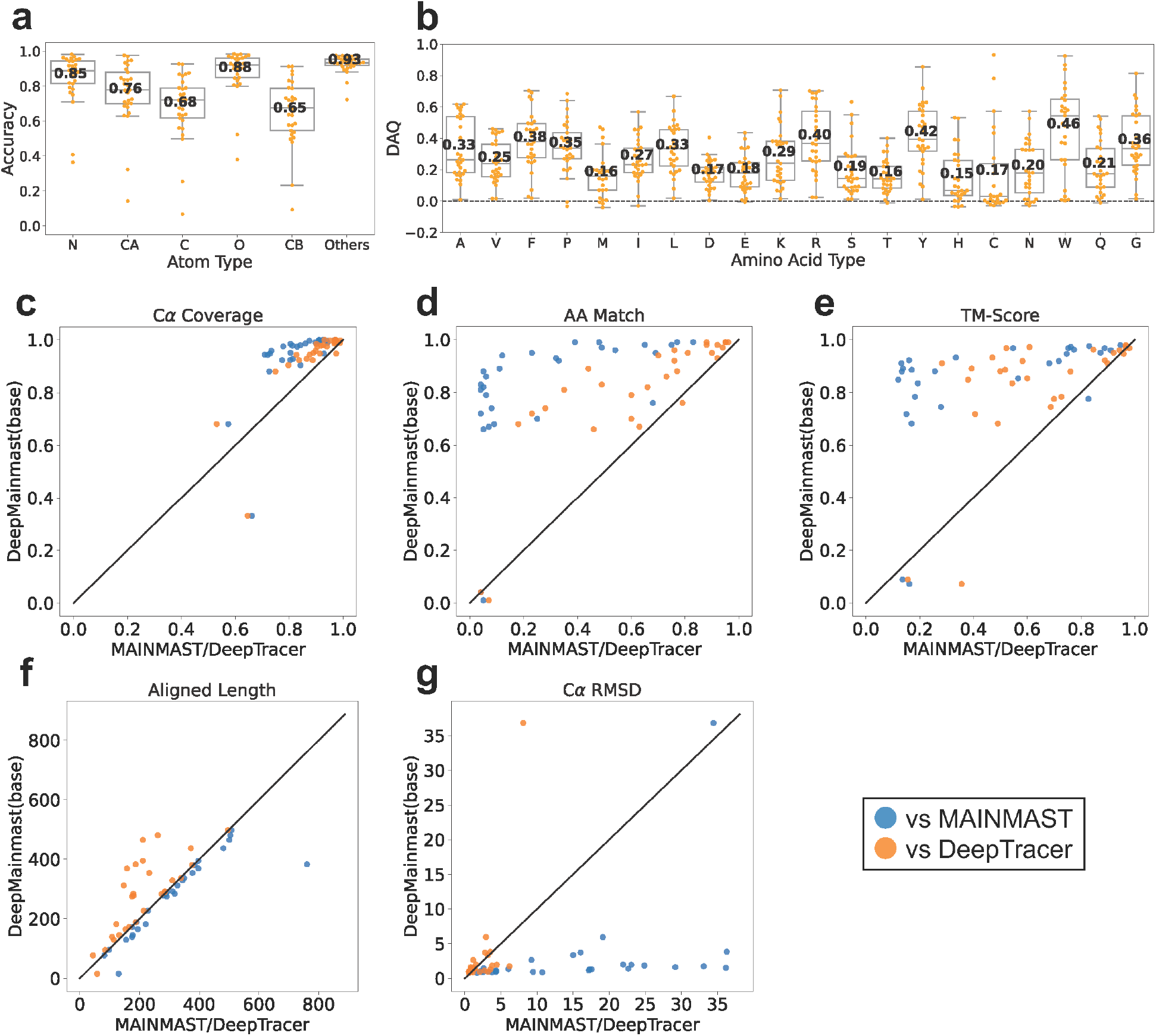
Sigle-chain modeling results on the 29 EM map dataset. **a**, the average atom detection accuracy for each map by Emap2sf. Each dot shows the value of individual map. The bold numbers shown are the average values across all the maps. In this box plot, the center line, the bottom and the ceiling in a box show the median, first quartile, and third quartile value, respectively. The boundaries of whiskers show 1.5 of the distance between upper and lower quartiles. An atom was considered to be correctly predicted if more than half of its corresponding grid points have correct atom assignment. **b**, The average DAQ(AA) scores of each map. A dashed horizontal line was drawn at DAQ = 0. **c**, the Cα coverage of the protein models. It is defined as the fraction of Cα atoms in a model that are placed within 3 Å to the correct position. The results by DeepMainmast(base) (the y-axis) were compared with MAINMAST (blue) and DeepTracer (orange) (the x-axis). **d,** the amino acid matching accuracy. It is defined as the fraction of Cα atoms in a model which are placed within 3 Å to the correct position and have the correct amino acid type assignment. **e,** TM-Score computed with the TM-align program. **f,** the length of aligned regions between the model and the native structure by TM-align. These regions were used to compute RMSD in panel g. **g**. Cα RMSD of protein models.

The first two panels in Fig. 2 show the accuracy of structure detection by the deep learning. Fig. 2a shows the atom detection accuracy. Atom positions were detected well with the average accuracy of five atom types ranging from 0.65 to 0.93. Of particular importance for the structure modeling protocol is the Cα atom detection, which achieved an accuracy of 0.76 on average for a map. The accuracy of detecting Cα atom was low, measuring below 0.4, for two maps, EMD-2364 and EMD-3246. The lower resolution of EMD-2364, at 4.4 Å, and the poor local map quality of EMD-3246 may have contributed to the low Cα atom accuracy. Fig. 2b shows DAQ scores of amino acids in the target proteins in their correct positions in the map, which is another critical information from deep learning because they were used to align protein sequences to detected Ca paths in the map. The DAQ score ranges from negative to positive values and a positive value indicates that the amino acid fits well into the position in the map. The overall positive DAQ scores in Fig. 2b indicate that Emap2sf successfully detected the correct position of amino acids in the map on average. The atom detection accuracy and DAQ scores of individual maps are provided in Supplementary Table 2 and Table 3, respectively.

The rest of the panels in Fig. 2 present the accuracy of structure models. Results of individual maps are provided in Supplementary Table 4. These panels correspond to the first-version models, which only include Cα atoms. Fig. 2c reports the Cα coverage, which is the fraction of Ca atoms in the native structure that were matched with any Ca atoms in the structure model within 3.0 Å. On average, DeepMainmast(base) identified 93.4% of Cα atom positions, which was higher than MAINMAST (82.5%) and DeepTracer (89.5%). Fig. 2d examines the accuracy of amino acid matching. It is the fraction of Cα atoms in the native structure that match with Cα atoms in the predicted model within 3.0 Å and also have the same amino acid type in the model. This metric is always lower than the Cα coverage in Fig. 2c because a match needs to also satisfy the amino acid type agreement. For computing matching Cα and matching amino acid type, we used the *phenix.chain_comparison* tool in the *Phenix* software^24^. The average amino acid match of the DeepMainmast model was 80.7%, while that of MAINMAST and DeepTracer models were 28.8% and 64.3%, respectively (Figure 2d). For almost all the targets, DeepMainmast(base) had a higher amino acid match than the other two methods.

The Cα coverage (Fig. 2c) and the amino acid match accuracy (Fig. 2d) are sequence order-independent metrics. On the other hand, the subsequent metrics in Fig. 2, TM-Score^25^ (Figs. 2e) and Cα RMSD (Fig. 2g) are sequence order-dependent metrics, which evaluate the topological accuracies of protein models. TM-Score ranges from 0 to 1, with 1 indicating the structures are identical. To compute the TM-Score for the DeepTracer models, we used a structural alignment program, TM-align^25^. DeepMainmast(base) clearly achieved a higher TM-score than MAINMAST and DeepTracer for all but 5 maps. The average TM-Score of DeepMainmast(base), MAINMAST, and DeepTracer were 0.83, 0.47, and 0.66, respectively. Compared to the structure models generated by DeepTracer, the models built by DeepMainmast(base) typically include more correctly placed amino acids, as indicated by the higher number of orange data points in Fig. 2f. In some cases, DeepTracer models contain Cα atoms that were not assigned to the correct sequence or position, causing them to be excluded from structural alignment by TM-align and resulting in shorter aligned lengths in Fig. 2f. In contrast, MAINMAST always includes all residues in the protein target, but DeepMainmast(base) occasionally misses a small number of residues (blue data points in Fig. 2f) due to its use of fragment-based modeling with the VRP solver. These missing residues can be filled in by Rosetta-CM in the subsequent step. In terms of Cα RMSD (Fig. 2g) the models by DeepMainmast(base) have lower value than DeepTracer models for 8 out of 29 cases (27.6%) despite that more residues were in general included in structure alignments considered. Against MAINMAST, DeepMainmast(base) had clearly better Cα RMSD results, with only one exception, EMD-2364. EMD-2364 (PDB: 4btg-A) is relatively at a low resolution of 4.3 Å and the protein is large, 761 residue-long. DeepMainmast identified Cα positions comparable to DeepTracer but was not able to assign correct sequence position. The Cα coverage by DeepMainmast(base) as 0.68, but the amino acid matching accuracy was 0.04.

In Supplementary Fig. 2, we examine the accuracy of full-atom models from DeepMainmast(base). The relative performance of the full-atom models to MAINMAST and DeepTracer (Supplementary Fig. 2) were essentially the same as what we observed in Fig. 2.

### Evaluation of single-chain modeling on another dataset of 178 maps

We further benchmarked DeepMainmast on a different dataset that contains 178 experimental maps at a resolution of 5 Å or better. This dataset was obtained from a recent work^9^ that proposed the CR-I-TASSER method. This time we examine models built by the full DeepMainmast protocol (Fig. 1), which includes integration of AF2 models. Four modeling methods were compared: (full) DeepMainmast, DeepMainmast(base) (i.e. without the integration of AF2), AF2, and CR-I-TASSER. For the two versions of DeepMainmast, we performed full-atom model building and refinement, then, the top-scoring model was selected using DAQ and DOT scores. CR-I-TASSER combines their protein structure prediction methods with prediction of Cα atom positions in an EM map using deep learning. This architecture is similar to the DeepMainmast protocol, which integrates DeepMainmast(base) with AF2, a structure prediction method. Since only TM-Score values of the CR-I-TASSER models were reported in their paper, we used TM-Score between a structure model and the native structure as the metric for this benchmark set. Results for individual maps are provided in Supplementary Table 5.

The first two panels in Figure 3 compares DeepMainmast(base) with CR-I-TASSER and AF2. Among 178 maps, DeepMainmast(base) had a higher TM-Score for 101 cases (56.7%) than CR-I-TASSER (Fig. 3a). When compared with AF2, DeepMainmast(base) had higher TM-Score for a slightly more than half, 91 cases (51.1%). When AF2 was integrated (Fig. 3c and 3d), DeepMainmast clearly outperformed the other two methods. Higher score was observed for 155 cases (87.1%) over CR-I-TASSER and 157 cases (88.2%) over AF2. Fig. 3e compares CR-I-TASSER with AF2. AF2 produced higher TM-Score models than CR-I-TASSER for 59.6% of the cases despite that AF2 does not consider EM map information.

**Figure 3.**
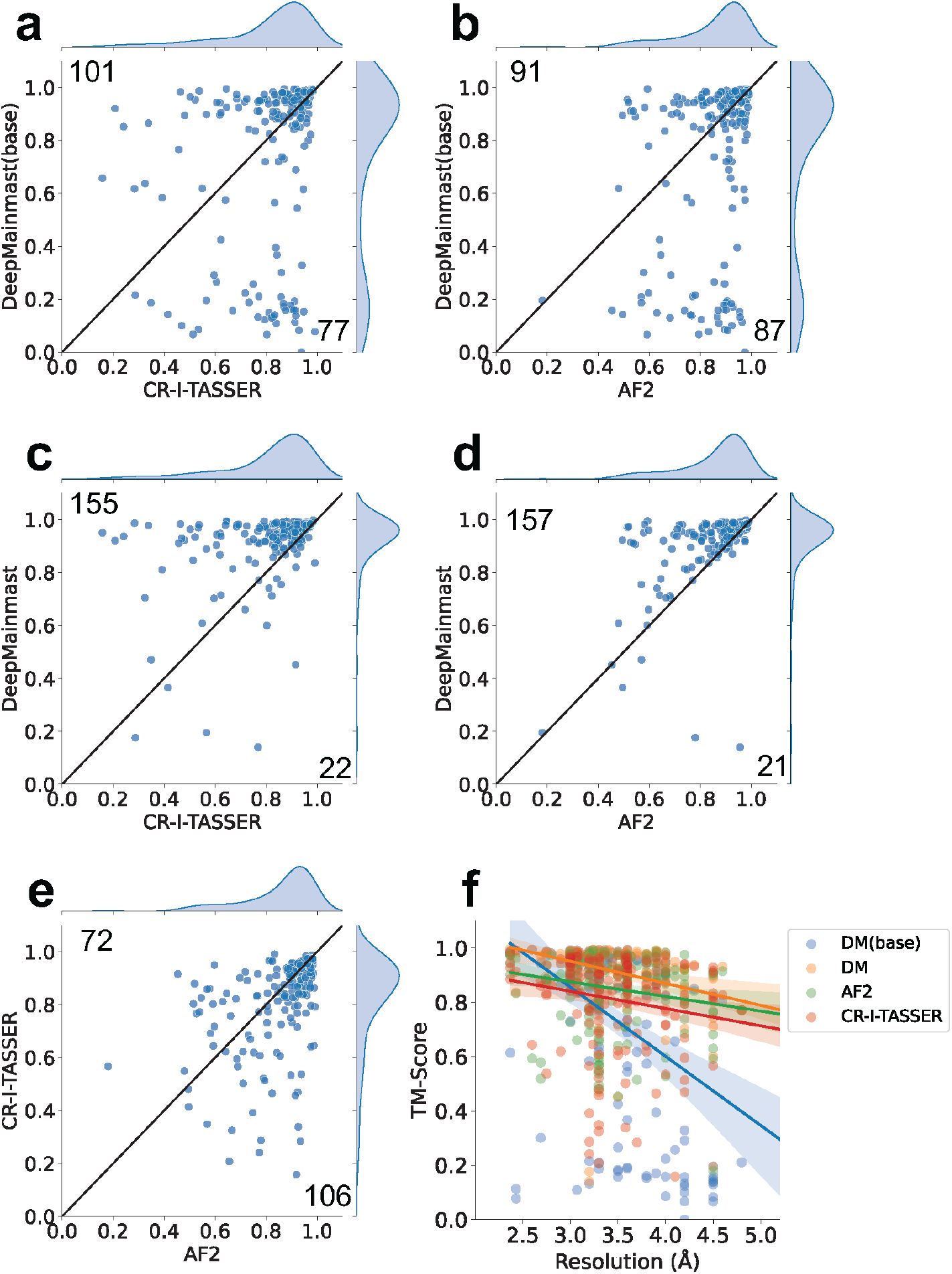
Single-chain modeling results on the 178 experimental maps. Panel a to e compare TM-Score of structure models generated by DeepMainmast, DeepMainmast(base) and two existing methods. The diagonal line in a plot represents the equal TM-Scores. The number at the top left and bottom right corners in a plot are the number of targets plotted above and below the diagonal line, respectively. **a,** comparison between full atom models of DeepMainmast(base) and CR-I-TASSER. **b,** DeepMainmast(base) and AF2. **c,** DeepMainmast and CR-I-TASSER. **d,** DeepMainmast and AF2. **e,** CR-I-TASSER and AF2. **f,** TM-Score of models relative to the map resolution. Blue, DeepMainmast(base); orange, DeepMainmast; green, AF2; red, CR-I-TASSER. The solid lines are regression lines. A colored area represents a 95% confidence interval.

Comparing marginal TM-Score distributions of DeepMainmast(base) and DeepMainmast (e.g. Fig. 3a and Fig. 3c), we noticed that there were a groups of maps that did not perform well by DeepMainmast(base), with a TM-Score of around 0.2 exhibiting a small peak. It turned out that these are maps of a low-resolution, lower than 4 Å. In Fig. 3f, we analyzed TM-Score of models relative to the map resolution. The figure shows TM-Score of models built by DeepMainmast(base) is negatively correlated with the map resolution, and they decrease particularly when the resolution becomes lower than 4 Å. At a resolution of 4 Å lower, map density starts to lose atom and residue level information, making it more difficult to correctly detect atom and amino acid residue information even with deep learning^2^. The other three methods in Fig. 3f do not show a decline in TM-Score as the resolution decreases because they use structure prediction methods that do not rely on map information. Therefore, in the full DeepMainmast protocol, AF2 compensates well for the density-based main-chain tracing performed by DeepMainmast(base).

To summarize, DeepMainmast(base) performed better than CR-I-TASSER and AF2 in terms of the number of models with higher TM-Score (Fig. 3a, 3b) and the margin has substantially increased when full DeepMainmast was considered. Particularly, DeepMainmast outperformed AF2.

### Examples of single-chain protein structure modeling

In Fig. 4, we present six examples of protein structure models built by DeepMainmast. The first three examples (Fig. 4 a-c) were from the dataset of 29 single-chain targets, which are shown to discuss the Cα tracing accuracy. Fig 4a is a 504 residue-long viral protein determined at a 2.8 Å resolution (PDB 5FOJ chain B, EMD-3246). For this map, Cα positions were detected well by DeepMainmast(base) and DeepTracer with high Cα coverages of over 0.90 by both methods. However, TM-Score of the two methods turned out to be largely different, with a score of 0.95 for the DeepMainmast(base) model and 0.47 for the DeepTracer model, due to the difference in the accuracy of assigning amino acid type and the sequence order of the protein to identified Cα positions. The application of RosettaCM made a small improvement of the DeepMainmast(base) from 0.93 to 0.95. Fig. 4b and 4c are examples from maps determined at a lower resolution, 3.5 Å and 3.8 Å, respectively. The TM-Score of the two methods were lower than the first example (Fig. 4a) for both methods, but DeepMainmast(base) maintained the score at 0.90 or higher. In the example of Figure 4c, a relatively large improvement of TM-Score from 0.85 to 0.90 was observed in the DeepMainmast(base) model by applying RosettaCM. This is due to the supplementation of missing residues by RosettaCM in the model including a loop of 11 residues and 12 other small gaps of one to four residues. Although the topology of the 11-residue-long loop was incorrect, other small fillings certainly contributed to the increase of TM-Score.

**Fig. 4.**
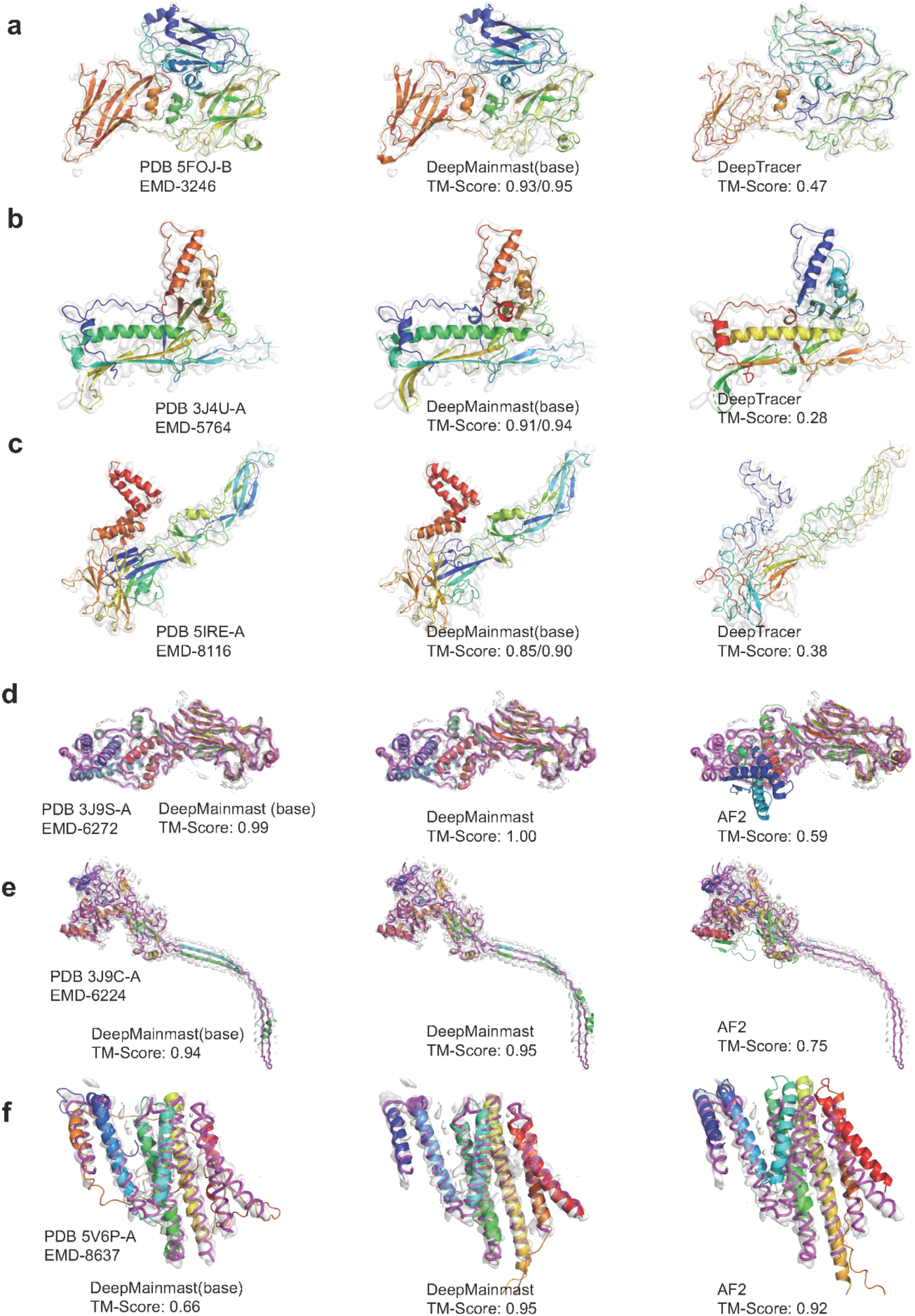
Modeling examples of single-chain targets. The first three panels, a-c are examples for discussing Cα tracing of DeepMainmast(base). Left, the native structure in PDB; middle, the DeepMainmast(base) model; right, the model by DeepTracer shown as reference. Chains were colored from blue to red to indicate the chain direction from the N-to C-terminus. Maps are shown at the author-recommended contour level. For DeepMainamast(base) models, TM-Score values of the Cα only model and the full-atom model after applying RosettaCM are shown on the left and right of ‘/’. **a,** RNA polyprotein of Grapevine fanleaf virus (PDB: 5FOJ chain B, EMD-3246, resolution, 2.8 Å). **b,** the major capsid protein of Bordetella phage BPP-1 (PDB 3J4U-A, EMD-5764, res. 3.5 Å). **c,** E protein of Zika virus (PDB 5IRE-A, EMD-8116, res. 3.8 Å). Panel d-f compares DeepMainmast(base), DeepMainmast, with AF2 and CR-I-TASSER. Models represented in cartoon are superimposed with the native structure in magenta. **d,** intermediate capsid protein VP6 (PDB 3J9S-A, EMD-6272, res. 2.6 Å). **e,** protective antigen pore protein (PDB 3J9C-A, EMD-6244, res. 2.9 Å). **f**, ubiquitin-protein ligase HRD1 (PDB 5V6P-A, EMD-8637, res. 4.1 Å).

The latter three examples in Fig. 4 are from the dataset of 178 maps discussed in Fig. 3. These models are to compare the full DeepMainmast with AF2 and also with CR-I-TASSER as the dataset was originally used in the work^9^ of CR-I-TASSER. Fig. 4d shows a 398-residue-long two domain structure of a virus capsid protein, VP6. The TM-Score of the AF2 model was 0.59 (Fig. 4d right), as the orientation of the two domains was incorrect although individual domain structures were folded correctly. This is a typical problem of AF2 when applied to a protein in a cryo-EM map. CR-I-TASSER also did not model the entire structure correctly, resulting in a TM-Score of 0.64 as it was reported in their work^9^. In contrast, DeepMainmast(base) was able to trace the almost entire main-chain correctly (TM-Score: 0.99), which was perfected to 1.0 by considering fragments from AF2 in the full DeepMainmast pipeline. The next protein (Fig. 4e) is a challenging target because it has a long, extended stem region of a β-sheet and the density around the stem is locally in a lower resolution and close to the density from other chains. CR-I-TASSER and AF2 built the correct fold for the globular domain of this protein but did not build the stem structure, resulting in TM-Score of 0.62 and 0.75, respectively. DeepMainmast(base) was able to trace the overall main-chain conformation including the stem, except for the hairpin loop at the tip of the stem because the map does not have clear density there. The hairpin loop was not recovered by using AF2 fragments either because AF2 did not model the region correctly. The last example (Fig. 4f) is a 407-residue-long protein that has a helix bundle fold. Since the EM map has a relatively low resolution of 4.1 Å, DeepMainmast(base) failed to build some α-helices and made wrong chain connections (Fig. 4f, left). AF2, on the other hand, built an overall accurate topology, but the relative positions of α-helices are slightly shifted, particularly the C-terminal helix (colored in red). DeepMainmast generated the most accurate model because it considered the local structure fragments from the AF2 model, which the DeepMainmast(base) could not build, and supplemented the density tracing result. This example illustrates how the AF2 model can enhance the modeling accuracy for challenging targets.

In Supplemental Figure 3 we analyzed how the model accuracy improved at three major steps in the DeepMainmast protocol. We used the three protein targets shown in Fig. 4d, e, and f for the illustration. Models were analyzed after the steps of Assembling Cα Fragments, Combining Models, and Building Full-Atom Models & Refinement in Fig. 1. In the first two targets, 3J9S-A and 3J9C-A, TM-Score distribution has progressively shifted to higher values. When models from DeepMainmast(base) (red points) and models that used AF2 fragments (black points) were compared, we can see that the AF2 fragments have contributed to producing high TM-Score models at the first fragment assembly step (blue boxes). But it is interesting to observe that DeepMainmast(base) models (black points) reached to equally high TM-Scores as models with AF2 fragments (red points) at the end of the three steps (green boxes). In the last target, 5V6P-A (Supplementary Figure 3c), DeepMainmast generated more accurate models than DeepMainmast(base) at all three steps. This is due to the low resolution of the map, and DeepMainmast(base) could not generate accurate Cα fragments in some regions as shown in Fig. 4f.

### Multi-chain structure modeling

We evaluate protein complex structure models built from cryo-EM maps, using 20 experimental maps of multiple chain complexes that were previously included in the maps of the 29 single-chain dataset used in Fig. 2. When we used these maps for testing the single-chain modeling ability, we segmented the maps to extract a single-chain region, but now we performed full-chain modeling using the entire maps. The number of chains in the maps ranged from 3 to 14 chains. The largest target, EMD-5764, has five chains with a total of 4,082 residues. For reference, we ran DeepTracer on their webserver with the same input data. Modeling results of individual maps are provided in Supplementary Table 6.

We first compare DeepMainmast with and without applying the chain ID assignment step (Fig. 5a). In all but one case, the chain ID assignment either improved or kept the TM-Score of the multi-chain complex models. As shown, the chain ID assignment algorithm can often substantially improve TM-Score by correcting fragment combinations for forming individual chains. In seven cases, TM-Score approximately doubled by the chain ID assignment.

**Fig. 5.**
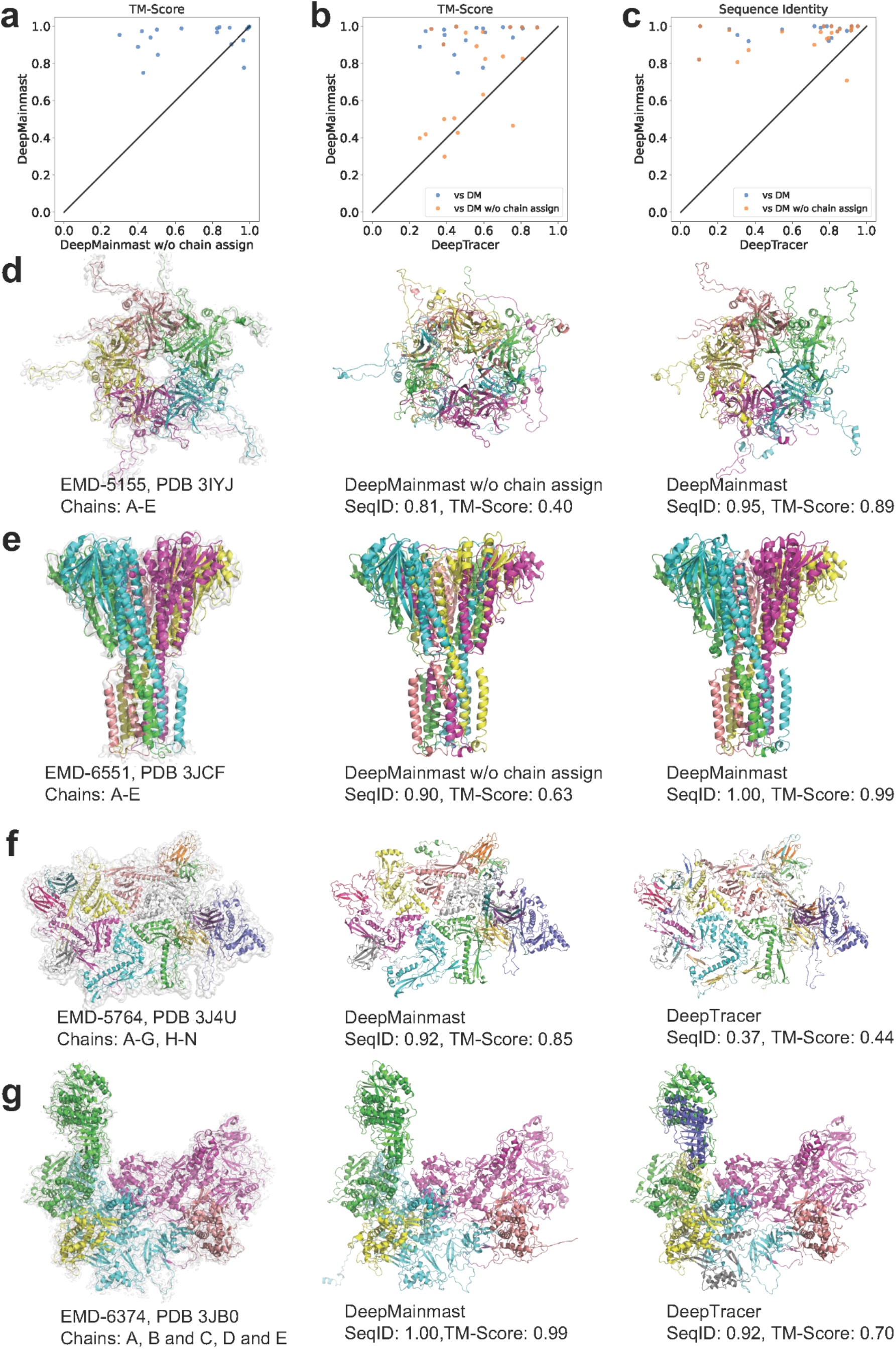
Modeling results of 20 multi-chain protein complex targets. **a**. Comparison of TM-Score of full-atom models of DeepMainmast with and without chain assignment. **b**. Comparison with DeepTracer in terms of TM-Score. Full-atom models of DeepMainmast with chain assignment (DM, blue) and without chain assignment (DM w/o chain assign, orange) were compared with DeepTracer models. **c.** Comparison with DeepTracer in terms of the sequence identity at aligned positions. TM-Score and the sequence identities were computed by MMalign^26^. **d and e.** The native structure and modeling results of DeepMainmast with and without the chain assignment. **d,** EMD-5155. PDB ID: 3IYI; 5 chains; resolution: 4.1 Å. **e,** EMD-6551. PDB ID: 3JCF; 5 chains; res.: 3.8 Å. **f and g.** The native structure and modeling results of DeepMainmast and DeepTracer. **f,** EMD-1461. PDB ID: 1QHD; 3 chains; res.: 3.8 Å. All three chains have an identical sequence. **g,** EMD-5764. PDB ID: 3J4U; 14 chains; res.: 3.5 Å. Chains A to G have an identical sequence and chains H to N have an identical sequence. Chains in the complex models were colored according to the chain ID assignment result. The chain color in a model show correspondence with the chains in the native structure.

**Fig. 6.**
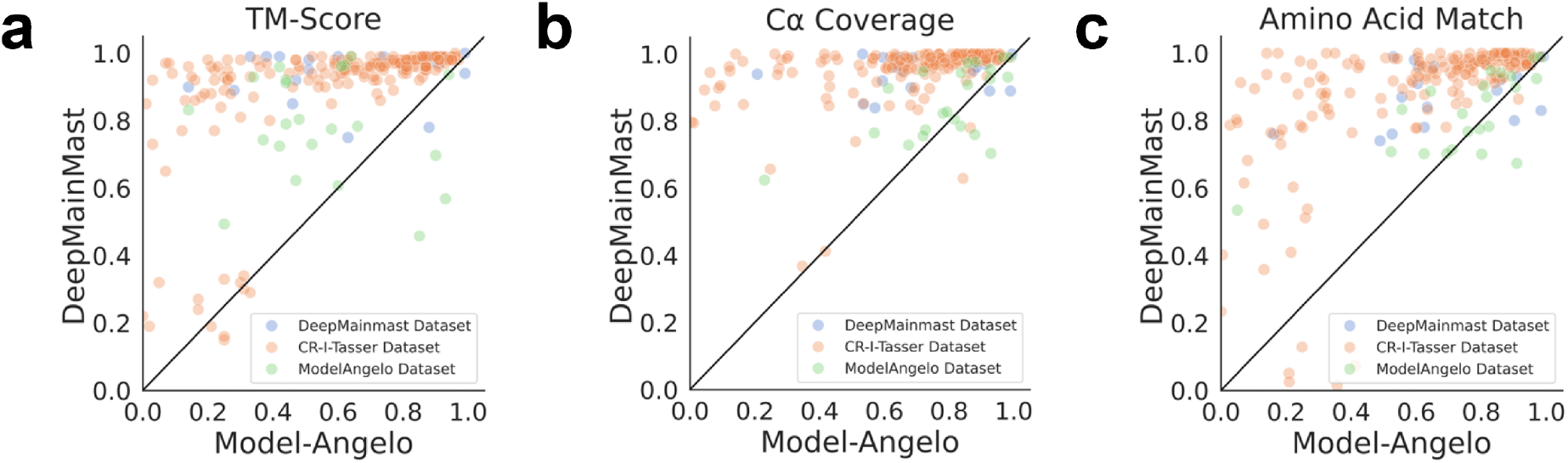
Modeling results on 3 datasets compared against ModelAngelo. The three datasets used are the 20 multi-chain targets used in Fig 5; the 178 single chain targets in the CR-I-TASSER dataset, and 22 multi-chain targets from the Model Angero paper. **a.** TM-Score comparison (DeepMainmast: 0.94, 0.90, 0.78 (overall 0.89), ModelAngelo: 0.64, 0.59, 0.55 (overall 0.59)). **b**. Cα coverage Comparison (DeepMainmast: 0.95, 0.96, 0.85 (overall 0.94), ModelAngelo: 0.73, 0.69, 0.80 (overall 0.71)). **c.** amino acid match (AA Match) comparison (DeepMainmast: 0.89, 0.89, 0.82 (overall 0.88), ModelAngelo: 0.71, 0.62, 0.76 (overall 0.64)).

Next, we compare the performance of five methods: DeepMainmast(base) and DeepMainmast with and without Chain ID assignment (four combinations) and DeepTracer using four metrics (Supplementary Table 7). Among the five methods, the full DeepMainmast protocol, including the chain ID assignment, showed the highest performance for all the metrics. In terms of the average TM-Score, DeepMainmast had the highest score of 0.94, followed by DeepMainmast(base) and DeepMainmast without chain ID assignment, which had a score of 0.74, DeepMainmast(base) without chain ID assignment (0.65), and DeepTracer (0.55), in this order. The scores of the all the metrics consistently improved when the chain ID assignment was applied in both DeepMainmast(base) and DeepMainmast. DeepTracer assigned Cα atoms well with a Cα coverage of 0.91. However, it had lower values for the other metrics compared with the other methods mainly due to a low amino acid matching and sequence alignment accuracy. Fig. 5b and 5c compare TM-Score and the sequence assignment accuracy of DeepMainmast and DeepTracer. DeepMainmast (data points in blue) had higher values than DeepTracer for all the 20 targets.

Fig 5d and 5e are for illustrating how the chain ID assignment improves complex models in the DeepMainmast protocol. Both examples, EMD-5155 (Fig. 5d) and EMD-6551 (Fig. 5e), are homo five-chain complexes. In both cases, DeepMainmast achieved high Cα coverage (0.81 and 0.94, respectively) and the sequence identity (0.81 and 0.90, respectively) in the models even without chain ID assignment (the middle panel), indicating that amino acid positions and types were well detected and modelled (Supplementary Table 6). However, TM-Score of the models were relatively low, 0.40 and 0.63 respectively in Fig. 5d and 5e, because local segments were swapped between chains. This problem was fixed by the chain ID assignment (right panels), resulting substantial increase of TM-Score to 0.89 and 0.99, respectively.

In Fig. 5f and 5g, we compare modeling results of DeepMainmast with DeepTracer. Fig. 5f is a homo trimer complex of Rotavirus VP6 protein (EMD-1461). For this target, DeepMainmast assigned the sequence well with a sequence identity of 0.90 without the chain assignment step, but the TM-Score was 0.63 due to fragment swaps (the middle panel). The chain ID assignment corrected the fragment combinations, improving the TM-Score to 0.99 (the right panel). The DeepTracer model had a TM-Score of 0.56 and the sequence identity of 0.55. Fig. 5f is a 14-chain complex of Bordetella phage BPP-1 (EMD-5764), which consists of seven homo-oligomer of major capsid protein (chain A-G) and seven cementing proteins (chain H-N). This is apparently a challenging target with many homo-oligomer chains and the intricate complex structure. DeepTracer had a Cα coverage of 0.88, slightly higher than DeepMainmast (0.85), but the overall TM-Score became low (0.44) due to less than half (0.44) amino acid matching accuracy and several incorrect chain connections. DeepMainmast also had a similar level of TM-Score of 0.51 without the chain ID assignment step, but the step substantially corrected chain assignment and recovered TM-Score to 0.85.

### Comparison with Model Angelo

We further conducted a comprehensive evaluation of the protein single chain and complex structure modeling performance on three distinct datasets from the papers of DeepMainmast, CR-I-TASSER, and Model Angelo^27^. The evaluation results are in Supplementary Table 8, 9, 10, respectively. DeepMainmast exhibited average TM-Scores of 0.94, 0.90, and 0.78, in the three test datasets, respectively, while Model Angelo had average TM-Scores of 0.64, 0.59, and 0.55, respectively. The average AA match for DeepMainmast was 0.89, 0.89, 0.82 on the three datasets, respectively, whereas Model Angelo’s AA match values were 0.71, 0.62, and 0.76, respectively.

## Discussion

We developed DeepMainmast, a deep learning-based method for *de novo* protein structure modeling for cryo-EM maps. Possible main-chain regions in the EM map are traced by connecting detected Cα positions by deep neural network, and they are aligned with target protein sequences using the scoring function, DAQ, which considers detected amino acid types along the trace. The entire modeling is considered as an optimization problem of fragment selections and mapping solved by the CP solver. Notable features of the DeepMainmast algorithm include the use of the VRP solver, which is well suited for modeling multiple chains, chain ID assignment correction for homo multimer targets, and the integration of AF2 models. Correct chain assignment for a homo multimeric complex is not a trivial task, and this is the first time the solution is implemented in a cryo-EM modeling tool. The integration of AF2 is a realistic solution for achieving high accuracy, which has complementary strengths to the density tracing from the map. The modeling is fully automated, and no human intervention is needed. Overall, DeepMainmast achieved better performance than AF2 and the current state-of-art *de novo* modeling methods we compared against. We believe DeepMainmast will be a useful and powerful tool for the structure biology researchers who use cryo-EM.

## Acknowledgments

This work was partly supported by the National Institutes of Health (R01GM133840 and 3R01GM133840-02S1) and the National Science Foundation (CMMI1825941, MCB1925643, IIS2211598, DMS2151678, DBI2146026, and DBI2003635).

## Author contributions

DK conceived the study. GT designed and implemented DeepMainmast and overall modeling protocol. XW coded and trained deep neural network and computed probability values of structure features for cryo-EM maps. GT and XW constructed datasets. GT, XW, and TN performed the computation and GT, DK, XW, and TN analyzed the data. DP participated in preparing codes to release and developed the CodeOcean pages and the Google Colab notebooks. GT and XW drafted the manuscript and DK edited it. All the authors read and approved on the manuscript.

## Data and code availability

The source code of DeepMainmast is made available at https://github.com/kiharalab/DeepMainMast. The free server is available at https://em.kiharalab.org/algorithm/DeepMainMast. It can run on Google Colab notebook https://github.com/kiharalab/DeepMainMast/blob/main/DeepMainMast.ipynb. We have also prepared capsules at CodeOcean at https://codeocean.com/capsule/9358532.

VESPER used in the pipeline is available at https://github.com/kiharalab/VESPER. DAQ is available at https://github.com/kiharalab/DAQ.

## Methods

### Detection of local structural properties in an EM map

The first step of DeepMainmast(base) is to detect local structural properties in an input cryo-EM map using deep learning. The deep learning method we use here, named Emap2sf (Emap to structural features), has a U-shaped Network (UNet3+)^20^ architecture with skip connections. The detailed architecture of 3D UNet3+ is shown in Supplementary Figure 1. For each grid point in the density map, Emap2sf computes probabilities of twenty amino acid types; and six atom types (N, Cα, C O, Cβ, Others). Emap2sf consists of two UNet3+ network architectures, one focused on amino acid type detection and the other for atom type detection. It takes density values extracted in a box of 32^3^ Å^3^ and outputs probability values for 20 amino acid types and 6 atom types at each grid in the boxes. Emap2sf is conceptually similar to our early work of deep-learning-based secondary structure detection, Emap2sec+^28^, but they have technical differences. The original Emap2sec+ uses a 3D Convolutional Neural Network (CNN) ResNet^29^ architecture and was trained on medium-to low-resolution (5 to 10 Å) cryo-EM maps. On the other hand, Emap2sf adopted a U-shaped Network (UNet3+) and was trained on maps with a higher resolution (2.5 to 5 Å) as this is the targeted resolution range for protein structure modeling by DeepMainmast(base).

### Training and validation datasets of experimental maps for Emap2sf

We first downloaded cryo-EM maps from EMDB^30^ that were determined at a resolution of 2.5 to 5.0 Å and have corresponding PDB entries deposited by the authors. Maps were removed if the deposited structures include DNA/RNA structures or unknown residues. To ensure the EM maps and associated PDB structures have proper alignments, we calculated the correlation between the experimental map and a simulated map from the deposited structure at the resolution of the experimental map. Maps were removed if the cross-correlation was lower than 0.65. Alignment between a map and the deposited structure was also manually inspected to confirm the agreement between the structure and the map. To build a non-redundant dataset, a map was removed if at least one protein chain pair from two maps share more than 25% sequence identity with each other. After applying all these steps, 237 cryo-EM maps remained. Among them, 197 maps (Supplementary Table 11) were used for training and validation of Emap2sf and 40 maps (Supplementary Table 12) were reserved as a test set.

The maps underwent pre-processing steps before fed into the neural network. First, the grid size was unified to 1.0 Å, if the original grid size is different, by trilinear interpolation. Second, density values in a map were normalized to [0.0,1.0] using minimum-maximum normalization, in the same way as our previous work^2^. Negative density values were set to 0, and 0 was used as the minimum value for a map. For the maximum value, we used the density value at the 98^th^ percentile and values larger than the cutoff were set to 1.

From each map, we collected boxes of a size of 32^3^ Å^3^ by scanning across the map along the three axes with a stride of 8 Å. We assigned each grid point in the box with an amino acid type and an atom type label that were taken from the closest heavy atoms of the closest residue located within 2.0 Å. If a grid point is close to more than two heavy atoms, then the closest heavy atom was used to provide the labels. If no heavy atoms were found within 2.0 Å of a grid point, the point was assigned as background. A box was discarded if less than 0.1% of the grid points within it had amino acid or atom assignment.

### Training the deep neural network of Emap2sf

Emap2sf has two separate UNets. The first UNet focuses on amino acid type detection and outputs probability values of twenty amino acids for each grid point to indicate the existence of corresponding amino acid. The second UNet is for atom type detection and outputs probability values for N, Cα, C O, Cβ, and Others. Both networks use the sigmoid activation function to output probability values ranging in [0.0, 1.0]. For each training batch, 64 density boxes were randomly selected from the 197 maps (164 for training and 33 for validation), which totaled around 210,000 and 45,000 boxes used in an epoch for training and validation, respectively. We used the dice loss^31^:

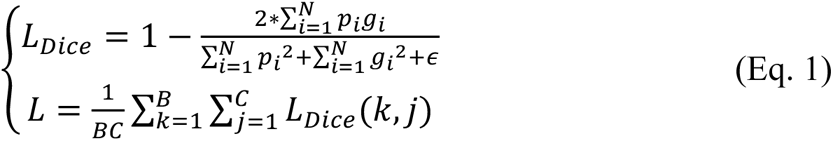

The first equation, *L_Dice_* represents the dice loss of a box P with computed probability values of amino acids/atoms at each grid point relative to the corresponding box *G* that contains correct assignments of amino acids/atoms. Here, *N* is the total number of grid points inside the box; *p_i_*_”_ ∈ P is the computed probability of the *i*-th grid point in the predicted box; *g*_”_ ∈ *G* is the binary ground truth of the *i*-th grid point, where 1 denotes the existing of such amino acid/atom in the grid point and 0 indicates no such amino acid/atom; ϵ is a smoothing factor with the value of 1e-6. *L* represents the overall loss of a batch of *B* examples with *C* class detection (amino acids/atoms); *L_Dice_*(*k*, *j*) represents the dice loss of *k*-th example’s *j*-th class detection. Deep supervision was used to apply the loss to the output of UNet from three different levels.

For the training of the network, we experimented with different combinations of a learning rate of [2e-5, 2e-4, 0.002, 0.02, 0.2] with an L2 regularization weight of [1e-6, 1e-5, 1e-4, 0.001, 0.01, 0.1] using the Adam optimizer^32^. Among these combinations, the learning rate 2e-3 with L2 regularization parameter of 1e-5 showed the best grid-wise Intersection-over-Union (IoU) of 27.6% for amino acid detection and 53.1% IoU for atom detection on the validation set, although the performances with different combinations were similar. We used the same hyper-parameters for training both atom type detection UNet and amino acid type detection UNet.

Training and validation of the network took about 5 days using around 255,000 data. The computation was on two paralleled NVIDIA TITAN RTX 24 GB GPUs with a NVLINK connection.

The network (Supplementary Fig. 1) includes three encoder blocks and two decoder blocks. Both encoder and decoder blocks are built upon Three-Dimensional Convolutional Layer (Conv3d), Batch Normalization Layer, and ReLU activation. The network has about 7.4 million parameters to train. In contrast, the training dataset has 210,000 boxes with 32^3^ values that need to be assigned 20 probability values for amino acids and 6 probability values for atom types. This amounts to over 137 billion values to predict. That comparison indicates that our model is not able to memorize the information to predict. Instead, it needs to learn a general pattern for accurate amino acid type and atom type detection.

Supplementary Table 2 shows the accuracy of Emap2sf on the 40-map testing dataset. To evaluate the detection of structure features by Emap2sf on an EM map, the ground truth label of amino acids and atoms was assigned to each grid point following the same rules to assign labels for training. On average, the grid-wise accuracy of amino acid detection is 36.9% and the accuracy of atom type detection is 72.2%.

### Tracing Cα paths

Using main-chain atoms detected in the cryo-E map by Emap2sf, Cα traces will be generated. Instead of connecting local points with a high density, we use the computed Cα probability of each grid points. As a preparation, representative points, called Local Dense Points (LDPs), in the EM map are generated from grid points with a Cα probability that is greater than a threshold Θ. Grid points with a low Cα probability is discarded. Then, LDPs are identified by the mean shift algorithm from the remaining points, which iteratively shifts each point toward points with a higher probability and cluster them locally to reduce points^5^.

Then, LDPs in the map are connected to produce Cα path(s) using the Vehicle Routing Problem (VRP) Solver^18^, which is a variation of the Traveling Salesman Problem (TSP) Solver. We used ORTools^18^ version 9.5.2237. Instead of using a single agent (salesman), multiple agents (vehicles) are used to visit and connect nodes (*x1*,..*xN*) in a graph, i.e. Cα LDPs in the map starting from a pseudo-node *x*0. The objective of VRP is to minimize the following cost function:

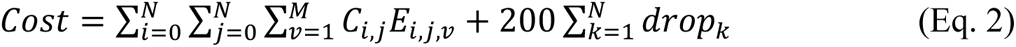

where *N* is the number of nodes (LDPs), *M* is the number of vehicles, *Ci,j* represents the cost of the path from the *i*-th node (*xi*) to *j*-th node (*xj*), *Ei,j,v* represents the edge and *Ei,j,v* = 1 if vehicle *v* directly visited from *xi* to *xj*; otherwise, *Ei,j,v* = 0. *dropk* controls the penalty cost if all vehicles did not visit the node *xk*. *dropk* = 1 if *xk* was not visited by any vehicles; otherwise, *dropk* = 0. All solutions of the path need to satisfy the following conditions:

All nodes need to be visited not more than once, i.e.

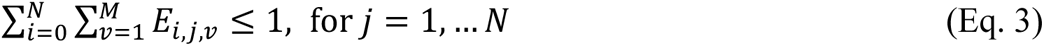

All paths (vehicles) must start from the pseudo-node *x*0 and connect to other node or stay at *x*0:

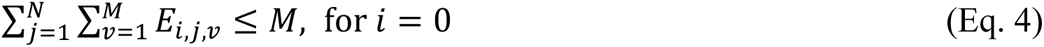

When a vehicle visits a node, the vehicle needs to depart from the same node:

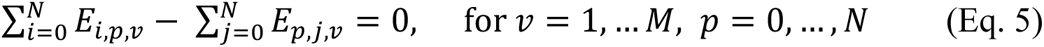

To define the cost *Ci,j* between two nodes (xi and xj), we introduced two ideas, (1) the closer the distance between two connected nodes to the ideal Cα-Cα distance (3.8Å^4,10^), the more likely the path is accurate, and (2) if Emap2sf does not predict a path between two nodes, then the two nodes likely do not have a connection between them. Following these ideas, we defined the cost *Ci,j* as:

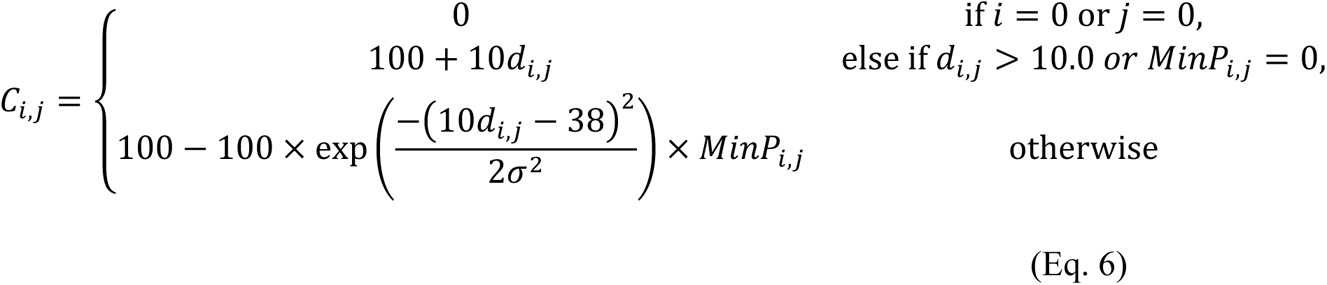

where *di,j* represents the Euclidian distance between *xi* and *xj*, *MinPi,j* is the lowest probability of backbone atoms (N, Cα, and C) on the straight line from *xi* to *xj*, σ is the standard deviation of the Gaussian function. The cost between *xi* and *xj* (*Ci,j*) will be close to zero if *di,j* is close to 3.8Å and *MinPi,j* is 1.0.

We varied values in the two parameters, Θ = [0.3, 0.4, 0.5], and M = [1, 5, 10] for single-chain targets or M = [5, 10, 20, 40] for multi-chain targets, and generated Cα paths for each combination to construct a pool of paths.

### Assigning sequence to Cα traces to generate Cα fragments

The obtained Cα paths are aligned with the amino acid sequence using the Smith-Waterman Dynamic Programming (DP) algorithm. To quantify the match between an amino acid *aai*, the i-th amino acid in a sequence, with the node *pj* in the path traced in the map, we used the DAQ score of amino acid types^2^. The DAQ score was developed to assess likelihood that an amino acid exists at a designated position in an EM map to assess the quality of protein models built from the EM map^2^. For a pair of *aai* and *pj*, DAQ is computed as:

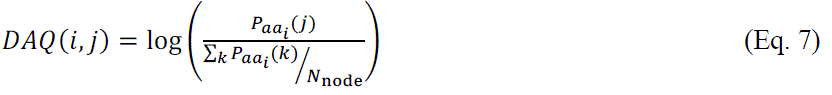

where P*_aai_*(*j*) is the computed probability of amino-acid type *aai* at the node *pj* by Emap2sf. The denominator is to normalize the score by the average value among all node *pk* in all Cα paths. *N*node is the total number of nodes in all the generated paths by the VRP Solver. Using DAQ score, a sequence-path alignment is computed with the following rule to fill a DP matrix *M*:

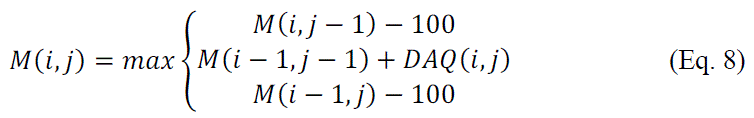

In the equation –100 is a large penalty for a gap in comparison with matching DAQ score, DAQ(i, j), which usually ranges between –2.0 to 2.0. For each Cα path, a total of four combinations of the threading process were performed, which come from two sequence directions and whether or not initializing negative DAQ with zero. The sequence threading on the Cα path was iterated Ψ times. For each iteration, we initialize the *DAQ*(*i*, *j*) with zero at aligned positions in the previous iteration round. By performing the iteration with the initialization of the *DAQ*(*i*, *j*), total of Ψ different sequence-path alignments were generated from the same Cα path. For single-chain and multi-chain targets, we used Ψ ∈ {5, 10} and Ψ ∈ {5, 10, 20}, respectively.

After the threading process, computed Cα fragments *fn* (*n*=1,…, *N*fragment) were evaluated by DAQ score. In the evaluation, we initialized negative *DAQ*(*i*, *j*) to zero. If the distance between the two connected nodes in the Cα fragment was larger than 7 Å, the Cα fragment was split into two parts. Small Cα fragments that have less than five nodes were removed. For a single-chain protein of about 100 to 900 amino acids, typically 1,000 to 50,000 Cα fragments are generated.

### Adding fragments from the AF2 model to the Cα fragment library

We use AF2 to predict the structure of the target protein and extract fragments from the AF2 model and use them in the subsequent modeling step only if they are locally consistent with the Cα fragments traced from the EM map. Among the five structure models that are generated by AF2, we use only one model that with the highest pLDDT score. The local consistency of the AF2 model is examined as follows^25,33^: First, nine residue-long fragments area extracted from the AF2 model and from the traced Cα fragments with a stride of one residue, and they are exhaustively compared. A pair of nine-residue-long fragments are superimposed and is considered as structurally consistent if the RMSD is less than 1.5 Å. Once a consistent pair is found, then the entire AF2 model is superimposed based on the alignment of the nine-residue-long fragments. Next, the alignment between the AF2 model and Cα fragments is updated by re-evaluating corresponding nodes between them, which should be located within a distance cutoff of 1.5 Å between each other. This process is iterated until the alignment converges or the number of iterations reached ten. If the length (the number of corresponding nodes) of the final alignment is less than the minimum alignment length Λ, the alignment is excluded. We used Λ ∈ {30, 50}. Then, the identified structurally consistent AF2 fragments are stored in the Cα fragment library. What frequently occurs is that a continuous segment with a loop structure flanked by secondary structures on both sides in the AF2 model aligns with two disconnected Cα fragments of the secondary structure regions. In such a case, if the gap (e.g. loop) in the alignment is less than 30 residues, the entire AF2 model region is stored in the fragment library in addition to the aligned regions in the model. Similarly, if an aligned region is close to the N-or C-terminus of the protein and the aligned region of the AF2 model is extended up to 10 residues toward the terminus and stored in the library. A small, isolated alignment region is removed from the final alignment if it is shorter than five residues.

There are four hyper-parameters used up to this point of the protocol. They are the threshold Θ (3 values) of the Cα probability, the number of vehicles M (3 and 4 values for single-and multi-chain targets, respectively), the number of different sequence alignment considered for a Cα path Ψ (2 and 3 values for single-and multi-chain targets, respectively), and the length of the fragment alignment to consider when matching with the AF2 model Λ (2 values). For each of the parameter combinations, we constructed a separate Cα fragment library, from which a Cα model is built in the subsequent step. The number of libraries for single-chain protein targets was 3 (Θ) x 3 (M) x 2 (Ψ) = 18, which do not include AF2 fragments and 3 (Θ) x 3 (M) x 2 (Ψ) x 2(Λ) = 36, which include AF2 fragments, thus, in total of 54 = 18 + 36. For multi-chain targets, the number of libraries was 3 (Θ) x 4 (M) x 3 (Ψ) = 36, which do not include AF2 fragments and was 3 (Θ) x 4 (M) x 3 (Ψ) x 2 (Λ) = 72 with AF2 fragments, thus 108 = 36 + 72 in total.

### Assembling Cα fragments using Constraint Programming (CP) Solver

For each Cα library, a Cα model is constructed by combining fragments with a CP Solver, which finds solutions for complex combination problems with constraints. Fragments are selected under the following constraints: Let *sn* (*n*=1, …, *N*fragment) be a DAQ score of each Cα fragment *fn*. The objective is to maximize

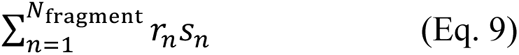

subject to

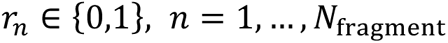

Under the constraint that prohibits the combination of two Cα fragments *fi* and *fj*, i.e.

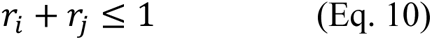

if *fi* and *fj* are an inconsistent pair, i.e. which is defined as a pair either (1) with a steric collision (< 2.5 Å for any Cα pair) or (2) with the same sequence position but placed at positions over 3.5 Å apart, or (3) with an inconsistent Cα− Cα separation between *fi* and *fj* that are placed at a distance larger than (the sequence separation) * 5 Å.

When *N*fragment is larger than 10,000, the CP Solver requires a large memory space and an extremely long computational time. To avoid a large computational cost for such a case, we split the Cα fragment library into randomly selected subsets with less than 10,000 fragments, and applied each subset separately and iteratively to the CP Solver.

### Fitting the AF2 models to the density map by VESPER

In addition to extracting fragments from the AF2 model, we also fit the entire AF2 model to the map using VESPER^19^, an EM map alignment program. We generated a simulated density map for the AF2 model at three different resolutions (5.0, 6.0, and 7.0 Å) using e2pdb2mrc.py^34^, and each resulting maps was superimposed onto the target EM map using VESPER. Ten best superimpositions were generated for each simulated map, which result in 30 superimpositions of the AF2 model on the EM map.

### Combining Assembled Cα fragment models

Assembled Cα fragment models undergo another round of fragment split and assembly process to combine different models. This process aims to fix potential problems, if any, such as gaps in the structure, imperfect overlaps of neighboring fragments, or errors in sequence assignments. At this point we have 108 Cα fragment-based models and 30 superimposed AF2 model to the map.

These models are classified into the following four groups: Group 1, models built by assembling Cα fragments obtained from tracing the EM map by DeepMainmast(base); Group2, models built from the extended Cα fragment libraries with those originating from DeepMainmast(base) and the AF2 model; Group3, models that are included in both Group 1 and Group 2; and Group 4: superimposed AF2 models to the EM density map by VESPER. For each of the four groups, models were cut into overlapping ten-residue-long fragments, from which a single Cα model is constructed by the CP Solver using the same condition and constraints. This process generates four models, one each from each group.

### Chain ID assignment for homo-oligomer targets

Assigning chain IDs in a multi-chain protein complex model is initially done during the sequence mapping step based on the assigned sequence. However, assigning chain IDs can be challenging for homo-oligomer cases, where different chains may have identical sequences. To ensure correct chain ID assignment for homo-oligomer targets, it is necessary to consider the consistency of assigned chain IDs in the structure model. This is particularly important for downstream applications that rely on the accurate identification of individual chains within the protein complex.

We run the CP solver with a specific objective function designed for optimizing chain assignment. If a Cα model of a homo-multimer has errors in the chain ID assignment, the chain ID assignment is swapped between equivalent local sequence regions of different chains (Supplementary Fig. 4a). Consequently, the chain models with the chain ID assignment error have a gap in their modelled structures, i.e., an unusually long distance between adjacent residues, which we aim to detect and correct. To start with, we define chain fragments along each chain model by checking the distance between adjacent residues. If adjacent amino acids are more than 10 Å apart at their Cα – Cα positions, we define the former and the latter parts from the gap as two separate chain fragments. Also, if residue numbers are discontinuous in a chain model, i.e. if a local region is missing from the model structure, the two parts are considered to be different chain fragments. The target function to optimize by the CP solver consists of two terms. The first term is to maximize the sum of DAQ scores from each chain fragment to ensure the overall good fit to the EM map. The second term is a penalty term to be minimized to resolve local structure inconsistencies between chains:

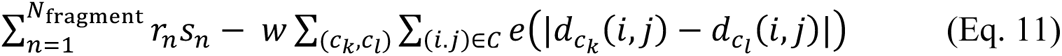

where *Nfragment* is the number of chain fragments in the homo-oligomer structure model, *sn* (*n*=1, …, *N*fragment) is DAQ score of the *n*-th fragment, and r_<_ ∈ {0,1}, *n* = 1, …, *N*_fragment_, which indicates inclusion or exclusion of the *n*-th fragment in the oligomer model. In the second term, *w* is a weight, set to *w =* 0.01, to balance the two terms in the target function. The first summation is to add up the penalty scores from every pair of chain combinations, *ck* and *cl*, and the second summation is to add up the penalty term *e* for every pair of amino acid residues, *i* and *j,* which satisfy the following condition, *C:* {(*i*, *j*)|(*i*, *j*) ∈ *C*common*AA*(*c*_*k*_, *c*_*l*_), *f*_*k*_(*i*) ≠ *f*_*l*_(*j*), *f*_*l*_(*i*) ≠ *f*_*l*_(*j*)}. *C*ommon*AA*(*c*_*k*_, *c*_*l*_) is a set of amino acid residue position pairs that exist both in chain *ck* and *cl*.

*f*_*l*_(*i*) is a function that tells the fragment ID *n* (1, …, *N*_fragment_) where the residue *i* of the chain *k* locates. Thus, the penalty term *e*(|*d*_*ck*_ (*i*, *j*) – *d*_*cl*_ (*i*, *j*)|) is considered only when two amino acids *i* and *j* locate in different chain fragments in both chain chain *ck* and *cl.* In the penalty term *e, d*_*ck*_ (*i*, *j*) is the Euclidian distance of residue *i* and *j* in the chain *ck. e* = 1 if *d*_*ck*_ (*i*, *j*) – *d*_*cl*_ (*i*, *j*)| > 2.0 Å, and 0 otherwise. The target function (Eq. 11) is optimized subject to

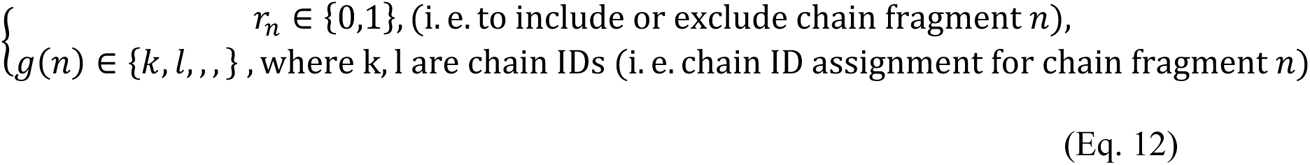

for *n* = 1, …, *N_fragment_*. *g(n)* is the function that assigns the chain fragment to a chain. See Supplementary Fig. 4b for illustration.

### Model evaluation by DAQ score and DOT score

We used the DAQ score^2^ and DOT score^19^ to evaluate the generated models. DAQ score (Eq. 6) is computed for all Cα positions in the full-atom model, and then the sum of the DAQ score is normalized by the number of residues in the model. DOT score is the objective function used in the map alignment program, VESPER. To compute the DOT score, we first computed a simulated density map from the full-atom model using e2pdb2mrc.py in the EMAN2 package at a 5 Å resolution. The grid spacing of the model’s simulated map and the target EM density map is converted to 2.0 Å, and then a unit vector at computed for each grid point *gi* (i=1, …, *Ngrid*). We use the density threshold value of 10.0 and 0.01 for the simulated map and the target EM map, respectively. In the map, the unit vector 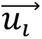 at *gi*, which points toward the local dense point, *yi* is defined as

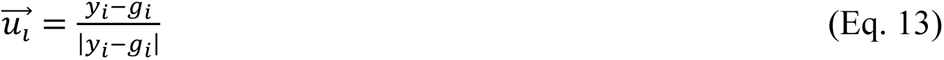

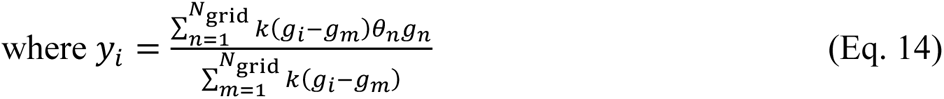

θ_<_ is the density value at the grid point *gn, k*(p) is a Gaussian kernel function that is defined as:

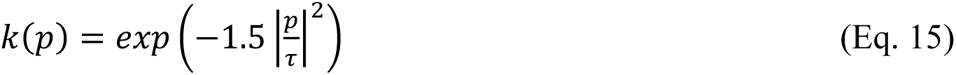

Where τ is a bandwidth and set to 8.0. Then, the DOT score, which is the sum of the agreement of unit vector pair, 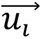 from the map and 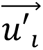 from the simulated map from the protein model is

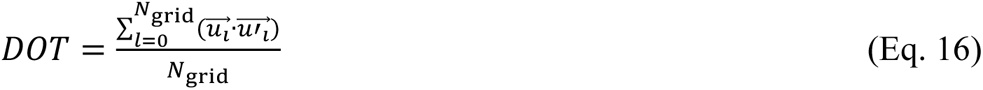

*N*grid is the number of grid points in the map.

The final score used to rank full-atom models is a simple sum of the normalized DAQ score and DOT score.

### Computational Time

We measured the computational time of the DeepMainmast protocol in Supplementary Fig. 5. We used a single GPU (Nvidia GeForce GTX 1080Ti, with 12GB memory) and four CPU threads on an Intel Xeon CPU E5-1650 v4. The computational time was split into three steps in the modeling protocol: up to Running deep learning and generating Cα fragments, up to Combining models, and until the end (Full-atom model building and refinement) (see the protocol in Fig. 1). The combining model steps takes the most time. It is generally proportional to the length of the protein but depends on the target protein structure. The required time is likely strongly influenced by the difficulty of modeling, i.e. more time is needed by the CP Solver for maps that need to explore a larger number of fragment combinations. On average, a single protein of up to ∼500 residues can be modelled within a few hours. Modeling of a multi-chain targets can be completed in 3-4 days. As the computational time was measured on a computer with a modest specification, the time can be substantially shortened if more CPU threads are used.

## Supplementary Information for

**Supplementary Figure 1.**
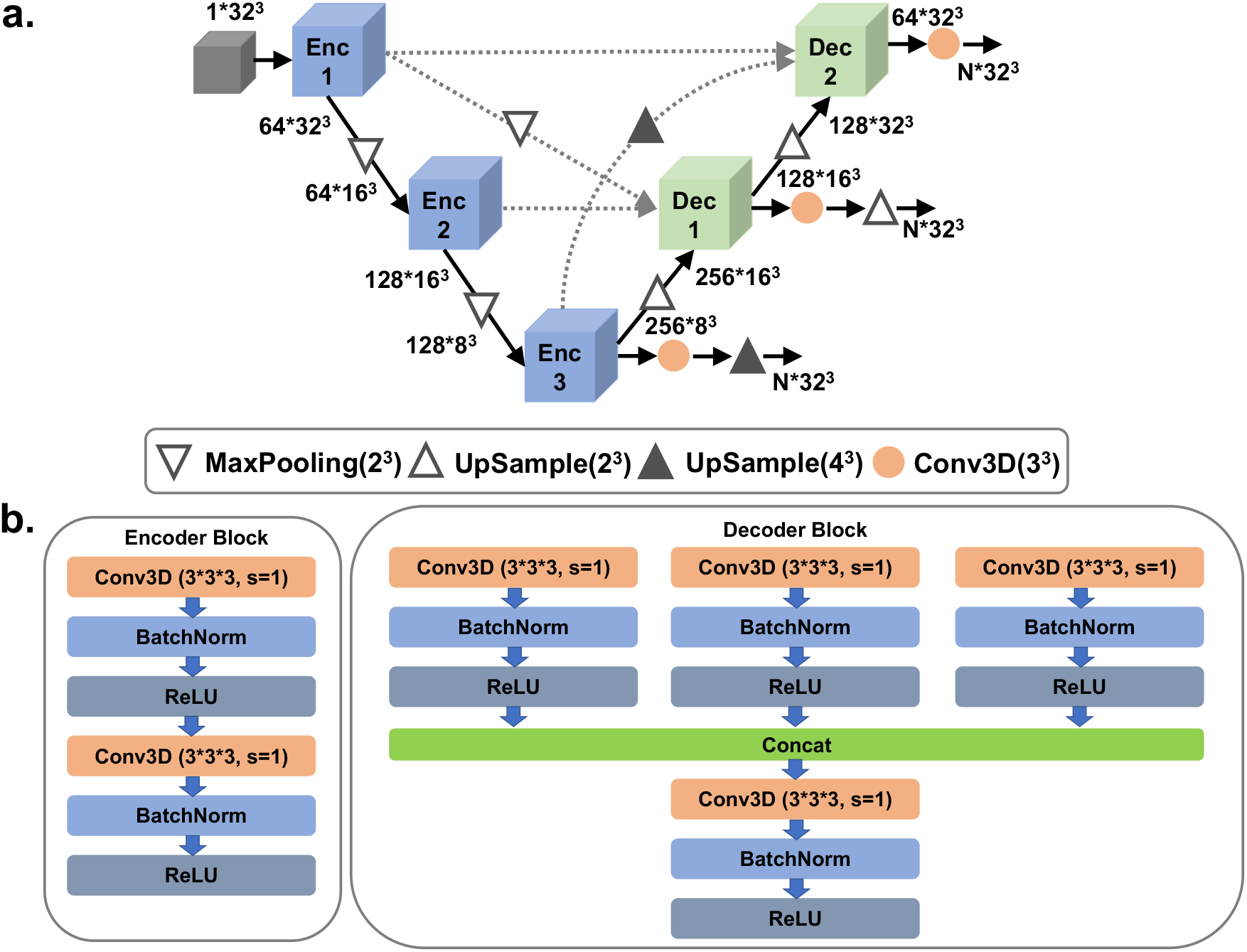
The network architecture of the deep learning method for local structure detection. The network architecture of Emap2sf (Emap to structural features), which is used to detect amino acid types, atom types, at each grid point in an input EM density map. **a.** the network architecture. The entire network is a 3D U-shape-based convolutional Network (UNet) with full-scale skip connections and deep supervisions. The numbers indicate the channel size of the corresponding layers. N is 20 for amino acid type detection UNet, N is 6 for atom type detection UNet. **b.** the encoder and the decoder blocks are shown. The encoder block (Enc in panel a; the decoder block (Dec in panel a). Conv3D, a 3-dimentional (3D) convolutional layer with the filter size of 3*3*3, stride 1 and padding 1. BatchNorm, a normalization layer that takes statistics in a batch to normalize the input data. ReLU, Rectified Linear Unit, a commonly used activation layer.

**Supplementary Table 1. The dataset of 29 single-chain proteins** (in a separate Excel file).

For each target entry, EMD-ID, PDB-ID, the map resolution, and the protein length are provided.

**Supplementary Table 2. The atom detection accuracy by Emap2sf for individual maps** (in a separate Excel file).

The atom detection accuracy for each map in the 29 single-chain protein dataset is provided.

**Supplementary Table 3. The average DAQ(AA) score of twenty amino acids of individual maps** (in a separate Excel file).

**Supplementary Table 4. Individual structure modeling results of maps in the 29 single-chain dataset** (in a separate Excel file).

For each map, Cα coverage, amino acid type match, TM-Score, Cα RMSD, and the number of modelled amino acids are provided. For DeepMainmast, two tables are provided. The first one is data from the Cα models, which correspond to Figure 2, and the second table is from full-atom models, which correspond to Supplementary Figure 2.

**Supplementary Figure 2.**
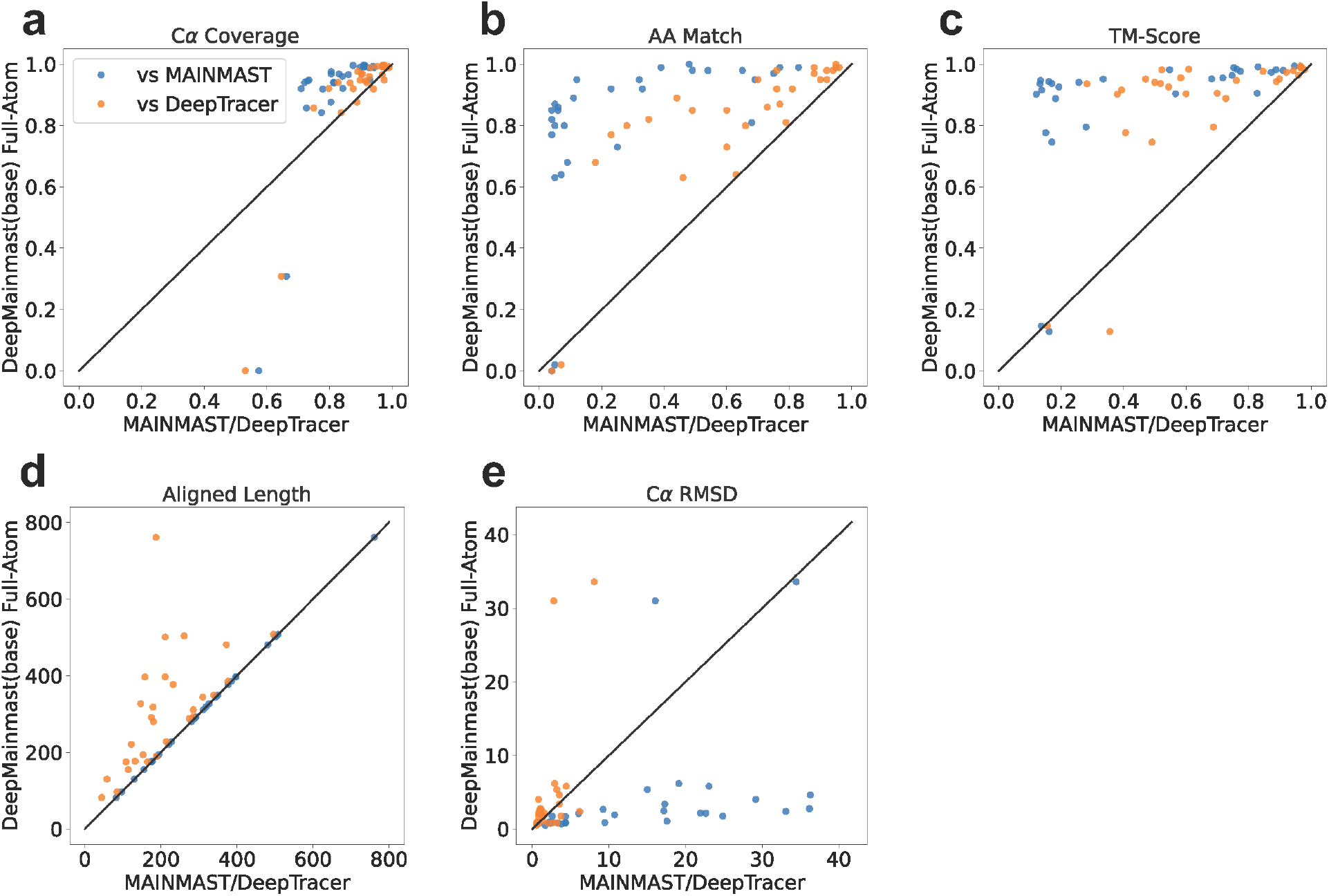
Modeling accuracy of full-atom models for the 29-map dataset. DeepMainmast(base) produced one Ca model for each target. For the Ca model, we ran Rosetta-CM, which fills missing regions (if any) and relax the structure, which produces 5 models. Out of them, we used the combination of the DOT score and the DAQ score, the same protocol as used in the final model selection step in the DeepMainmast pipeline (Fig. 1), to select the model for comparison. As for full-atom models for MAINMAST, following the MAINMAST protocol, MDFF was used to generate 500 full-atom models from one Ca model, among which the top-scoring full atom model was selected. **a**, the Ca coverage. **b**, the amino acid matching accuracy. **c**, TM-Score. **d**, the length of aligned regions between the model and the native structure by TM-align. These regions were used to compute RMSD in panel **e**, Cα RMSD of protein models.

**Supplementary Table 5. Individual structure modeling results for maps in the 178 single-chain dataset** (in a separate Excel file).

This dataset was obtained from the work of Zhang X et al (2022). (Zhang, X., Zhang, B., Freddolino, P. L. & Zhang, Y. CR-I-TASSER: assemble protein structures from cryo-EM density maps using deep convolutional neural networks. Nat Methods 19, 195-204, doi:10.1038/s41592-021-01389-9 (2022)). It includes 178 cryo-EM density maps of single-chain proteins at a resolution better than 5 Å. For each map, PDB ID, EMD-ID, map resolution, and TM-score by DeepMainmast(base), DeepMainmast, Alphafold2 (AF2), and CR-I-TASSER are provided. The results of CR-I-TASSER were obtained from the above-mentioned paper.

**Supplementary Figure 3.**
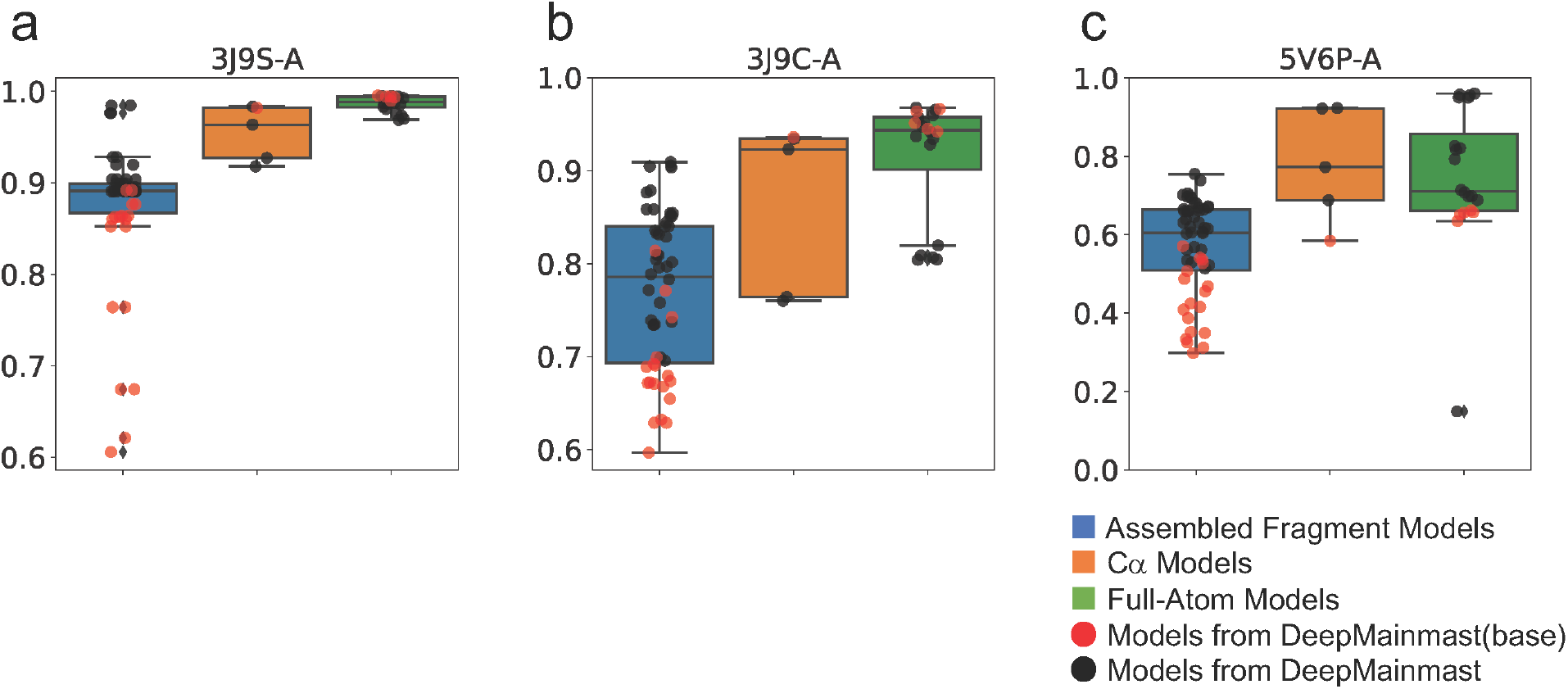
TM-Score distribution of models generated at major steps in the DeepMainmast protocol. For the three target proteins shown in Fig. 4 d, e, and f, all models generated at the three major steps were evaluated in terms of TM-score. **a.** Models for PDB 3J9S chain A (Fig. 4d). **b.** Models for PDB 3J9C chain A (Fig. 4e). **c.** Models for PDB 5V6P chain A (Fig. 4f). In each panel, blue, orange, and green boxplots show TM-Score distribution of the models generated by the “Assembling Ca Fragments”, “Combining Models,” and “Building Full-Atom Models & Refinement” steps, in Fig.1, respectively. In these three steps, 54, 4, and 20 models were respectively generated. Red circles represent the models generated by the DeepMainmast(base) protocol. Black circles represent the models additionally generated by DeepMainmast. In the box plots, the middle line in a box corresponds to the median, and the top and bottom ends of a box represent quartiles. The upper and lower whiskers represent 1.5 *the interquartile range. Black diamond represents the outlier from the whiskers.

**Supplementary Table 6. Individual structure modeling results for maps in the 20 multiple-chain dataset** (in a separate Excel file).

Results of four methods are shown. DeepMainmast(base), with and without applying the chain ID assignment; DeepMainmast with and without applying the chain ID assignment, and DeepTracer. The chain composition column shows the number of chains in the complex and chains that have identical sequences. Chain IDs connected by = have an identical sequence. Chains separated by: have different sequences.

Four results are shown for a target map by each method: Ca coverage; Amino acid matching accuracy; the sequence identity; which is defined as the sequence identity of the aligned regions between the model and the native structure by TM-align; and TM-Score. Plots in Fig. 4 a, b, c are based on these raw data.

**Supplementary Table 7.** Average accuracy values of the multi-chain targets.

|  | **Cα Coverage** | **AA match** | **TM-score** | **Sequence Identity** |
| --- | --- | --- | --- | --- |
| **DeepMainmast(base)** | 0.95 | 0.72 | 0.74 | 0.87 |
| **DeepMainmast(base) w/o chain assign** | 0.84 | 0.71 | 0.65 | 0.85 |
| **DeepMainmast** | **0.95** | **0.89** | **0.94** | **0.97** |
| **DeepMainmast w/o chain assign** | 0.93 | 0.87 | 0.74 | 0.94 |
| **DeepTracer** | 0.91 | 0.63 | 0.55 | 0.67 |

The average values from the 20 target maps. Individual map results are provided in Supplementary Table 6.

**Supplementary Table 8. Individual structure modeling results compared to model-angelo for 20 maps in DeepMainmast test dataset (in a separate Excel file).**

Results of DeepMainmast and model-angelo are shown. The chain composition column shows the number of chains in the complex and chains that have identical sequences. Chain IDs connected by = have an identical sequence. Chains separated by: have different sequences.

Four results are shown for a target map by each method: Ca coverage; Amino acid matching accuracy; the sequence identity; which is defined as the sequence identity of the aligned regions between the model and the native structure by TM-align; and TM-Score. Plots in Fig. 6 a, b, c are based on these raw data.

**Supplementary Table 9. Individual structure modeling results compared to model-angelo for 178 maps in CR-I-TASSER test dataset (in a separate Excel file).**

Results of DeepMainmast and model-angelo are shown. For each target entry, PDBID, ChainID are provided.

Four results are shown for a target map by each method: Ca coverage; Amino acid matching accuracy; the sequence identity; which is defined as the sequence identity of the aligned regions between the model and the native structure by TM-align; and TM-Score. Plots in Fig. 6 a, b, c are based on these raw data.

**Supplementary Table 10. Individual structure modeling results compared to model-angelo for 22 maps in model-angelo test dataset (in a separate Excel file).**

Results of DeepMainmast and model-angelo are shown. For each target entry, EMDID is provided.

Four results are shown for a target map by each method: Ca coverage; Amino acid matching accuracy; the sequence identity; which is defined as the sequence identity of the aligned regions between the model and the native structure by TM-align; and TM-Score. Plots in Fig. 6 a, b, c are based on these raw data.

**Supplementary Table 11. The training dataset of 197 maps** (in a separate Excel file).

For each target entry, EMD-ID, PDB-ID, and the map resolution are provided. These 197 maps were used for training and validation of Emap2sf.

**Supplementary Table 12. The test dataset of 40 maps** (in a separate Excel file).

For each target entry, EMD-ID, PDB-ID, and the map resolution are provided. These 40 maps were used as testing dataset of Emap2sf. The testing results were provided in Supplementary Table 2.

**Supplementary Figure 4.**
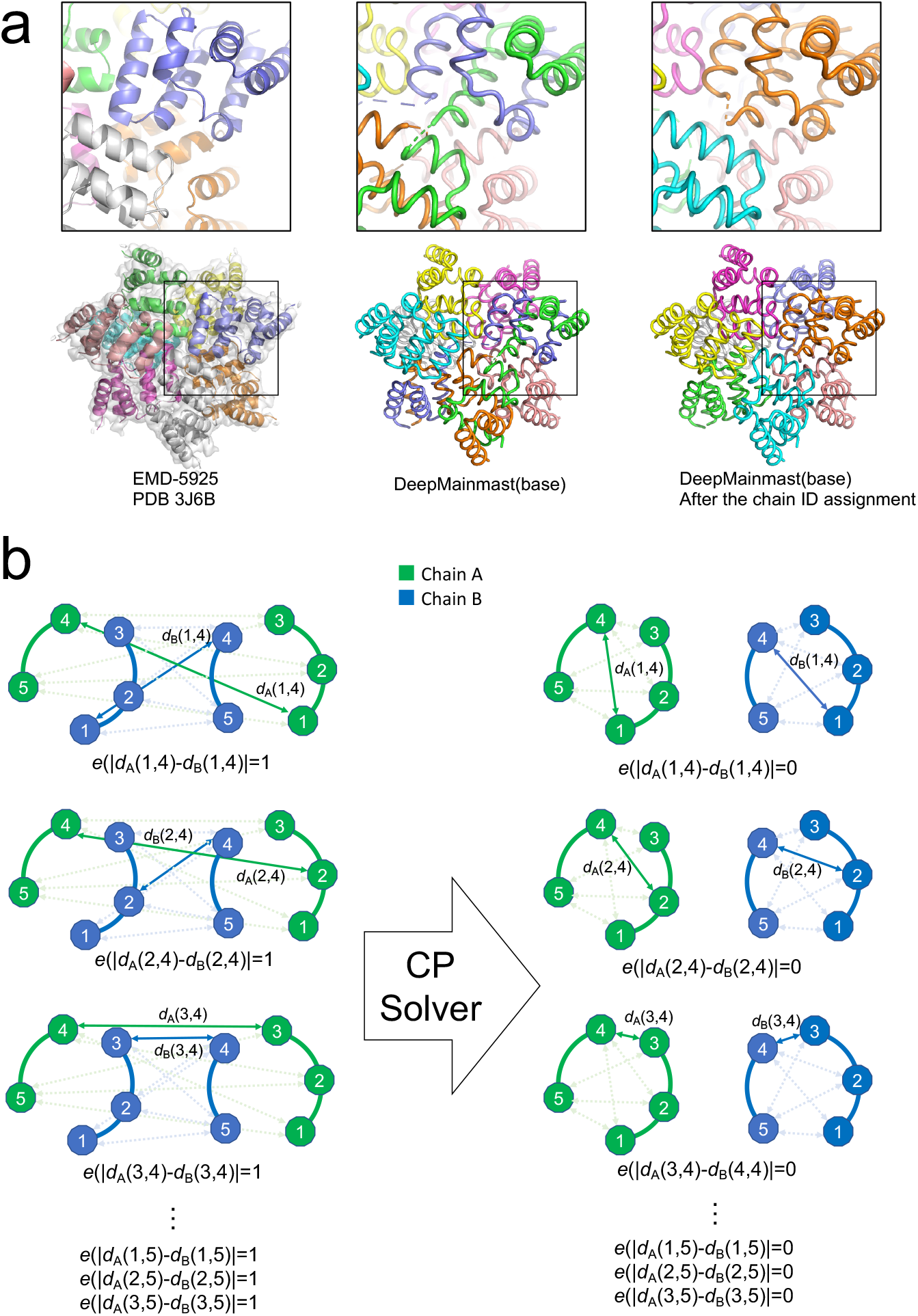
Chain ID assignment in the DeepMainmast protocol. **a**. Example of chain ID assignment for EMD-5925. The deposited model (PDB ID 3J6J) consists of homo octamer structure. All models were colored by chain ID. The magnified images highlight a region where different chains interact. The left column shows the deposited model (PDB 3J6J) of EMD-5925. The middle column shows the model generated by the DeepMainmast(base) protocol prior to the chain ID assignment step. The right column shows the DeepMainmast(base) model after the chain ID assignment step is completed. As shown, chains are correctly connected and identified. **b.** Illustration of the chain ID assignment for a homo-dimer target. In this example, two five-residue-long models with different chain IDs (green: chain A, and blue: chain B) are shown. Numbered circles represent amino acid residues and the number is the residue number in the sequence. For the chain ID assignment, DeepMainmast maximizes the object function (Eq.11) that consists of a DAQ score term and a penalty term. The penalty term is intended to have similar structures in different chains of homo-oligomers. On the left and right columns, we illustrated the computation of the penalty term *e*() before and after the chain ID assignment, respectively. Arrows between two residues with solid lines indicate the distance between residues *i* and *j* for chain A and B models (dA(*i*,*j*) and dB(*i*,*j*)). If the |dA(*i*,*j*) – dB(*i*,*j*)| > 2.0 Å, *e*() = 1. Since the models of chain A and B has different structures, all penalties *e*() is 1 before the chain ID assignment. After the chain ID assignment, all penalties *e*() were reduced to zero.

**Supplementary Figure 5.**
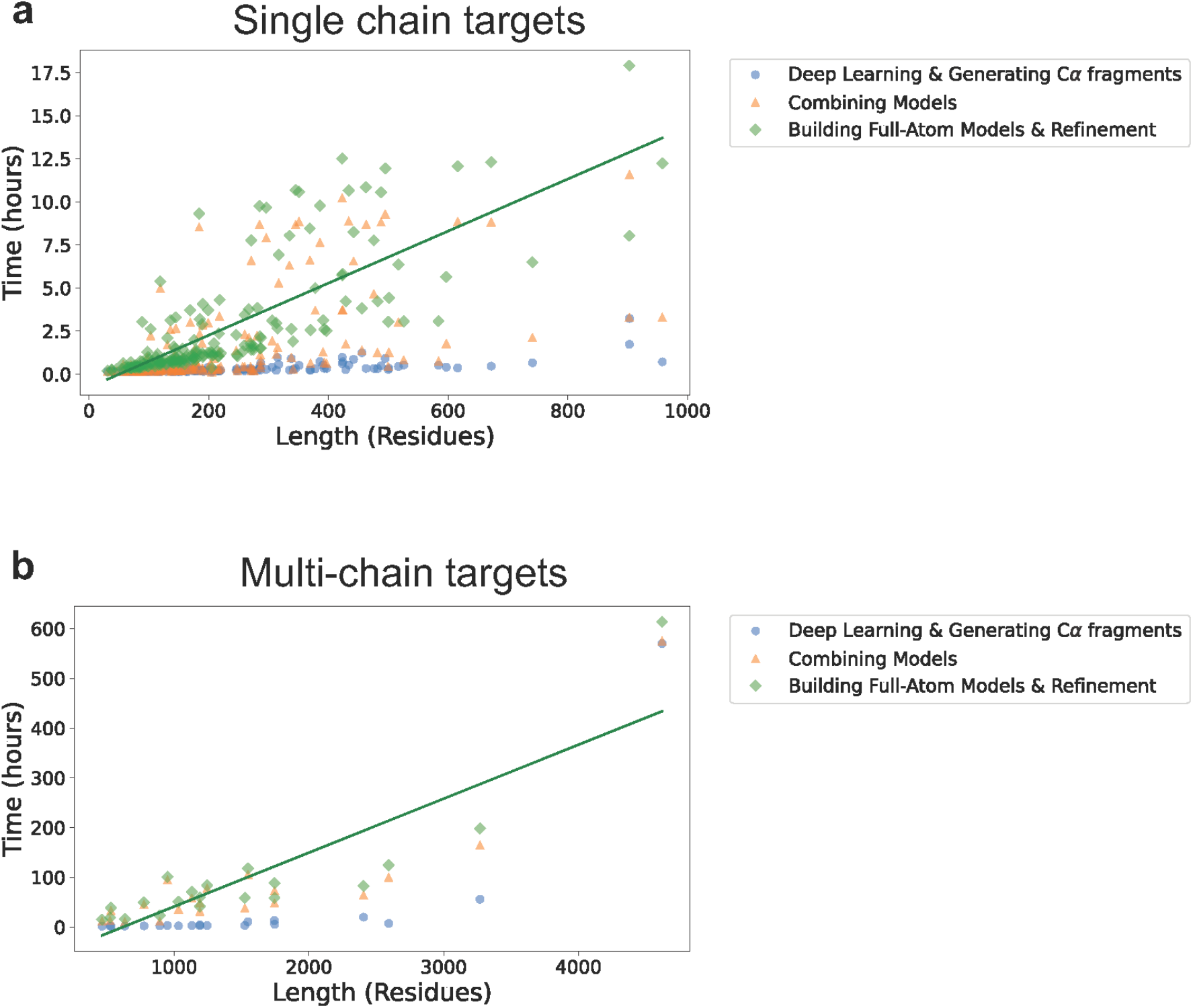
The computational time of DeepMainmast. **a**. Computational time of the DeepMainmast protocol on the dataset of 178 single-chain targets. **b.** Computational time of the DeepMainmast protocol on the datasaet of 20 multi-chain targets. For this experiment, we used one GPU card (Nvidia GeForce GTX 1080Ti, 12GB memory) and four threads on one CPU (Intel Xeon CPU E5-1650 v4). The plots show the computational time in three colors (blue, orange, and green), corresponding to the (1) the steps up to the “Assembling Cα Fragment”, (2) the steps up to the “Combining Models”, and (3) the steps up to “building Full-Atom Models & Refinement” in Fig.1, respectively. The green solid lines represent the regression lines of the total computational time.

